# NeuroImgAgent: a multi-agent platform for cross-species fMRI analysis with evidence-gated quality control

**DOI:** 10.64898/2026.11.19.767770

**Authors:** Jianyu Yuan, Hongtao Zhang, Liwen Wan, Yuanxi Shen, Zhifei Zhu, Yexin Han, Zeping Xie, Pingfu Wang, Qian Wan, Xiaojing Long, Chao Zou, Dong Liang, Hairong Zheng, Ye Li, Yi He

## Abstract

Cross-species functional MRI analysis requires coordinating specialized workflows and assessing intermediate outputs. We present NeuroImgAgent, a multi-agent platform driven by large language models that translates natural-language requests into workflows for humans, macaques, rats and mice. Quantitative checks and vision-language model assessments determine whether processing continues, enters bounded recovery or stops. Across 268 paired cases, preprocessing metrics were broadly comparable to those of fMRIPrep, AFNI and RABIES. With GPT-5.5, NeuroImgAgent corrected 111 of 120 induced orientation errors and passed all 52 intent-recognition and planning tasks, 8 of 9 recovery tasks and 11 of 14 tool-construction tasks on NeuroAgentBench. In 120 paired challenge cases, the full configuration increased usability from 76.7% to 92.5%, as assessed by blinded experts, and reduced unusable outputs delivered without warning from 18.3% to 4.2% of cases relative to quality-control ablation.

## Main

Functional MRI (fMRI) analysis depends on a chain of interdependent decisions spanning data organization, image preprocessing, confound handling and quality control. The Brain Imaging Data Structure (BIDS) standardizes dataset organization^1^, while tools and workflows such as fMRIPrep, AFNI and RABIES support reproducible processing in human, non-human-primate and rodent studies^2–4^. DeepPrep further illustrates how combining deep learning with workflow management can accelerate neuroimaging processing^5^. These advances standardize and automate many processing steps, but researchers still need to select species-appropriate workflows, adjust parameters when necessary and assess whether intermediate outputs are usable. These decisions can be challenging for researchers with limited neuroimaging or programming experience, particularly when analyses span different species and acquisition protocols.

Quality control (QC) often requires manual review, and undetected processing errors can compromise downstream analyses. Quantitative image-quality features can support automated assessment^6^, and studies of motion confounding show that preprocessing can leave systematic artefacts in functional-connectivity estimates^7,8^. Small-animal MRI adds challenges associated with anatomy, image scale and acquisition conditions^9^. Summary metrics can miss problems in image orientation, brain coverage, brain masks or registration, whereas manual visual inspection can be time-consuming and subjective. Orientation errors can escape detection unless spatial consistency is checked explicitly^10,11^. Vision–language models (VLMs) have been applied to biomedical image interpretation^12^. These advances motivate testing VLMs for QC using standardized MRI views alongside quantitative measures, with validation across species, error types and model configurations.

Large language model (LLM) agents offer a complementary approach to automation by translating requests into plans, selecting tools and updating their actions in response to observed outcomes^13,14^. Scientific agents have demonstrated these capabilities in chemistry^15^, single-cell analysis^16^ and biological design under human supervision^17^. GeneAgent uses external biological databases to verify generated interpretations^18^. An evaluation of LLM agents for atomic force microscopy found that strong performance in domain question answering did not guarantee reliable experimental execution. The agents also had difficulty coordinating tasks and were sensitive to small changes in prompting^19^. CopilotJ connects a conversational multi-agent system with ImageJ to support bioimage analysis and reusable workflows^20^. The AI Neuroscientist preprint demonstrated an LLM-based interface for QC, modelling and visualization of functional near-infrared spectroscopy data^21^.

Applying agents to cross-species fMRI requires careful assessment of intermediate spatial outputs. Species differ in image properties and reference spaces, and a completed processing step can still produce an inaccurate registration or brain mask. These challenges motivate an evidence-gated framework that uses quantitative and visual checks to guide execution and records the tools, parameters and evidence underlying its decisions. Recovery should be restricted to predefined repairs or registered fallbacks within retry limits. Detected fatal QC failures should block downstream processing if they remain unresolved. Evaluation should compare outputs with those of established workflows and assess output usability and the results of downstream analyses alongside task completion.

We developed NeuroImgAgent as a cross-species orchestration platform designed around these requirements. The platform profiles input data, identifies the species and translates natural-language requests into species-appropriate workflows for researcher approval before execution. A coordinator manages parallel execution across subjects, QC and evidence-linked reporting. Detected failures can trigger a recovery loop that diagnoses the problem and selects a predefined repair or registered fallback. The platform reruns only the affected branch and reassesses its outputs. Recovery attempts are limited, and downstream processing stops with an explicit warning if a detected fatal QC failure remains unresolved. We compared preprocessing outputs from NeuroImgAgent with those from fMRIPrep, AFNI and RABIES across human, macaque, rat and mouse datasets. NeuroAgentBench evaluated intent recognition, planning, fault recovery and tool construction. We also evaluated VLM-guided orientation correction and QC of registration and brain masks. Profiler and QC ablations assessed the roles of these components in species identification and output usability. Finally, we reanalysed the Midnight Scan Club dataset to examine whether NeuroImgAgent reproduced previously reported individual and individual-by-task patterns in functional-network similarity^22,23^.

## Results

### NeuroImgAgent integrates multi-agent orchestration with evidence-gated quality control

NeuroImgAgent translates natural-language requests and MRI inputs into reviewable fMRI workflows for humans, macaques, rats and mice (**Fig. 1a**). A coordinator organizes profiling, planning, execution, QC and reporting as a stateful graph (**Fig. 1b**). PROFILE inventories the available modalities and metadata and identifies the species; PLAN selects a compatible pipeline and validates its dependency graph. A human-in-the-loop (HITL) step allows researchers to revise and approve the plan before execution. EXECUTE then invokes registered neuroimaging tools and processes subjects in parallel, recording checkpoints to support recovery. QC evaluates node outputs, and REPORT links results to the corresponding processing and QC evidence (**Extended Data Fig. 1**). For eligible requests that require capabilities absent from the registry, the optional Toolsmith generates candidate tools that must pass validation before registration.

**Fig. 1:**
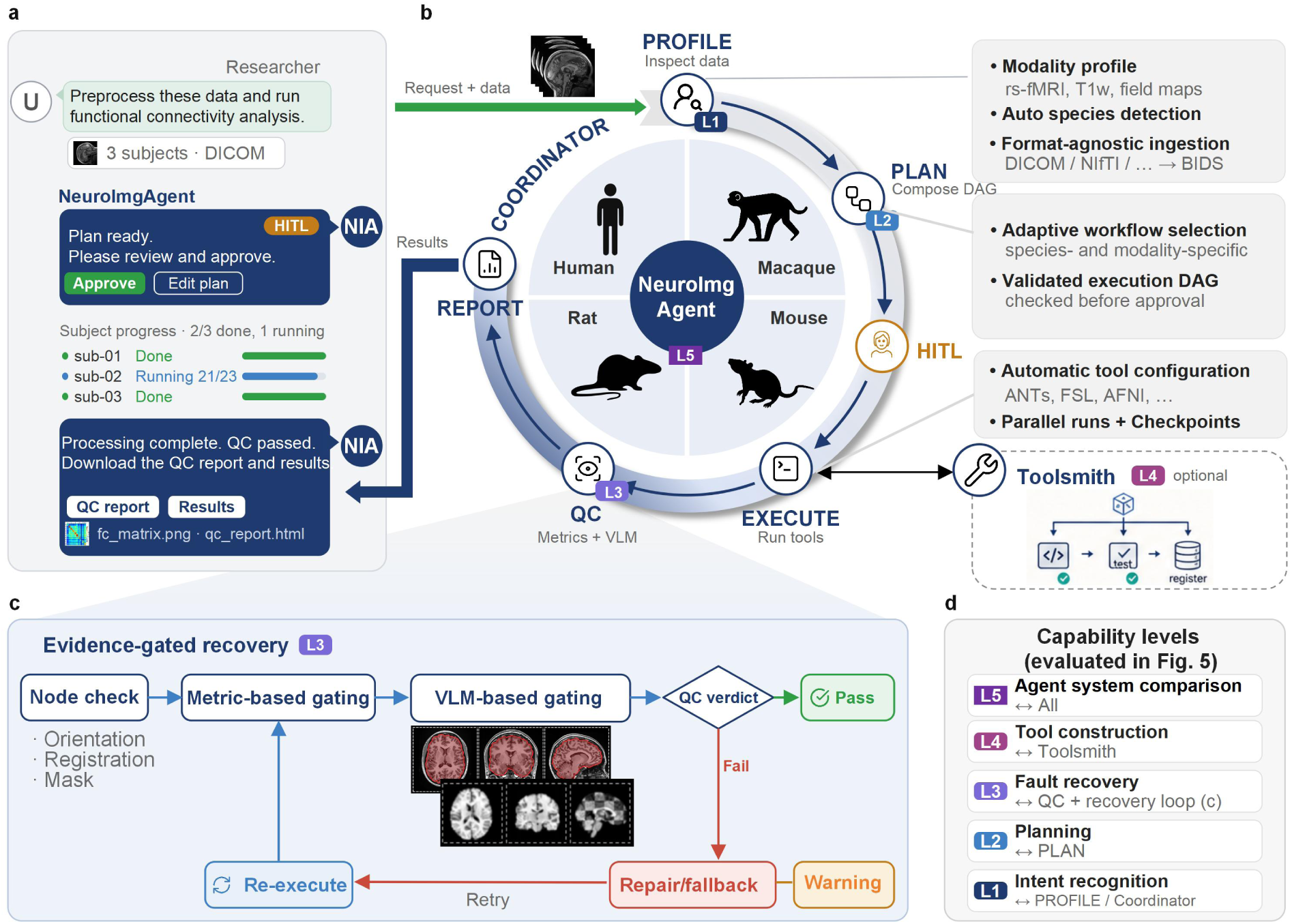
NeuroImgAgent: a multi-agent platform for cross-species fMRI analysis with evidence-gated quality control. **a**, User interaction and human-in-the-loop review. The researcher submits a natural-language request together with the raw data. NeuroImgAgent returns a proposed workflow for approval (human-in-the-loop, HITL), reports subject-level progress during execution, and delivers the processed data, the QC report and the final results. **b**, Coordinator-guided multi-agent workflow. A coordinator orchestrates five agents in sequence (arrows): PROFILE performs modality profiling, format-agnostic ingestion (DICOM/NIfTI → BIDS) and automatic species identification; PLAN selects a species- and modality-specific workflow and composes a validated execution directed acyclic graph (DAG), which is released only after human approval; EXECUTE configures tools automatically and runs subjects in parallel with checkpoints; an optional Toolsmith builds, tests and registers new tools into the toolbox for EXECUTE; QC evaluates every node with quantitative metrics and vision–language model (VLM) inspection; REPORT assembles evidence-linked outputs. The platform supports human, macaque, rat and mouse data. **c**, Evidence-gated recovery (expanded from QC in b). Each node (orientation, registration, brain mask) passes through metric-based gating followed by VLM-based visual gating. A PASS verdict lets execution continue; a FAIL verdict triggers diagnosis and repair or fallback, followed by re-execution and reassessment. **d**, Hierarchical agentic capability levels used for evaluation: L1 intent recognition (PROFILE/coordinator), L2 planning (PLAN), L3 fault recovery (QC and the recovery loop in c), L4 tool construction (Toolsmith) and L5 agent-system comparison (all agents; end-to-end analysis from raw data to report). Task-level performance at L1–L4 and the end-to-end L5 comparison with a single-agent baseline are quantified in Fig. 5. Coloured tags (L1–L5) mark the capability level exercised by each component.

QC uses file-integrity checks, task-specific measurements and standardized image views to determine whether execution can proceed between processing nodes (**Fig. 1c**). For image orientation, spatial registration and brain masking, deterministic checks are combined with VLM review according to task-specific QC rules. PASS and non-fatal WARN outcomes permit continued execution. Fatal FAIL outcomes block downstream processing. For failures eligible for recovery, structured diagnosis guides predefined parameter adjustments, upstream retuning or selection of a registered fallback. The affected node is then re-executed and reassessed. Retry limits and circuit breakers constrain recovery. Unresolved fatal QC failures stop downstream processing and are reported with an explicit warning. Model judgements, deterministic decisions and recovery histories are retained as distinct records.

We evaluated the platform using NeuroAgentBench, which separates intent recognition and routing (L1), data-aware planning (L2), fault diagnosis and bounded recovery (L3), tool construction and reuse (L4), and comparison of workflow execution with a single-agent system (L5; **Fig. 1d**). Task-based analysis and interactive visualization are illustrated with a human checkerboard experiment (**Extended Data Fig. 2**). Representative resting-state and task-based analyses are shown in **Supplementary Videos 1 and 2**, respectively.

### Cross-species preprocessing yields broadly comparable outputs

We compared NeuroImgAgent with fMRIPrep 24.1.1 for human data, an AFNI-based workflow for macaque data and RABIES 0.5.1 for rat and mouse data (**Fig. 2 and Supplementary Tables 1–3**). The paired benchmark comprised 170 human cases, 30 macaque runs, 30 rat cases and 38 mouse cases. Where required, inputs to the reference workflows were manually prepared by reorienting images, reorganizing files or providing an anatomical reference derived from echo-planar imaging (EPI) data for macaques without T1-weighted images. NeuroImgAgent processed the same scans without manual input preparation. Framewise displacement (FD) and temporal signal-to-noise ratio (tSNR) were summarized descriptively and showed broadly similar values between workflows within each species (**Fig. 2a,d,g,j**).

**Fig. 2:**
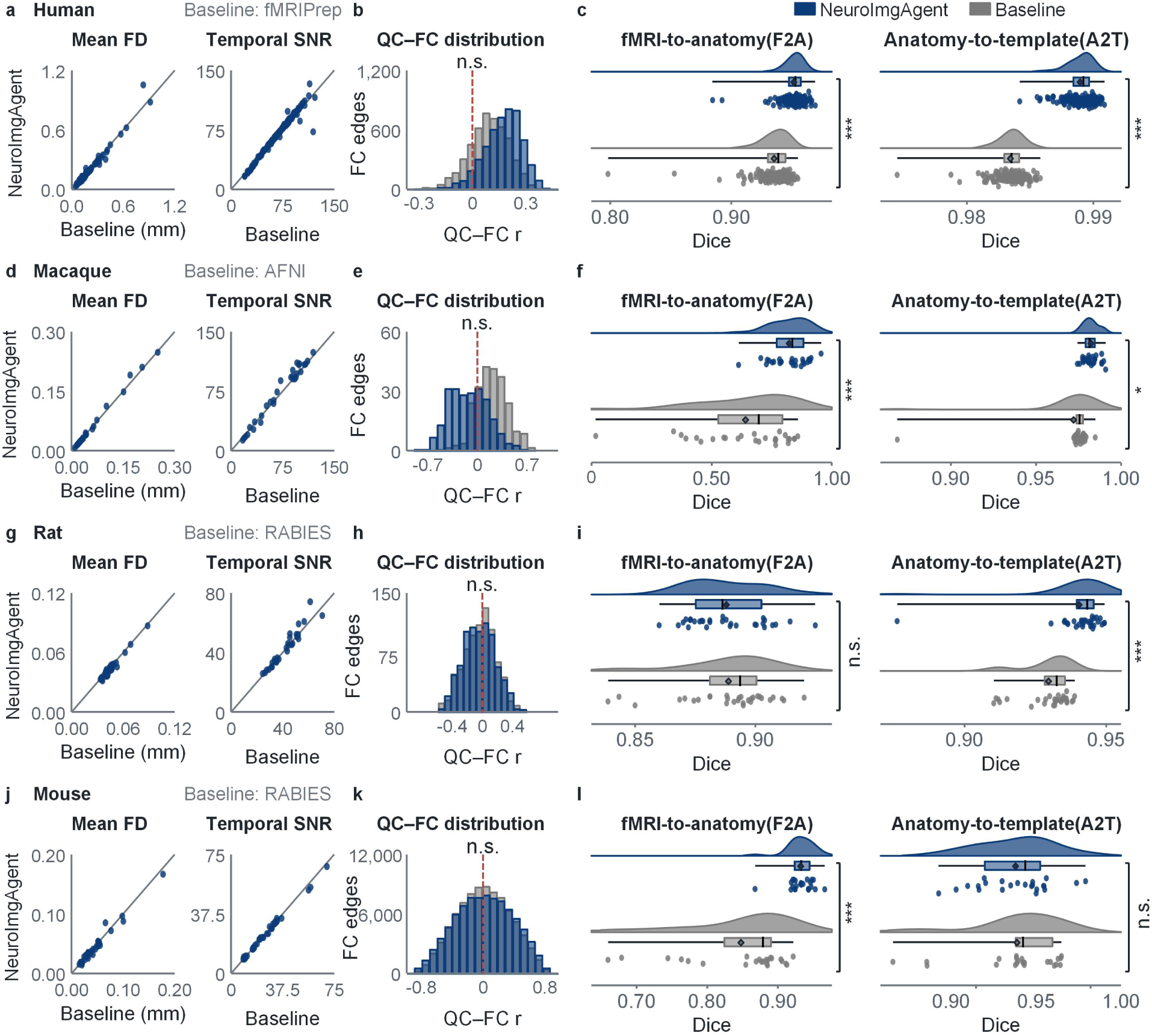
Cross-species performance comparison of NeuroImgAgent with established preprocessing pipelines. **a–c**, Comparison of preprocessing metrics between NeuroImgAgent and fMRIPrep on human data (n = 170 pairs). **a,** Agreement between NeuroImgAgent and fMRIPrep in mean framewise displacement (FD) and temporal signal-to-noise ratio (tSNR). **b**, Quality-control–functional-connectivity (QC–FC) correlations. The mean absolute correlation coefficient (|r|) was 0.170 for NeuroImgAgent and 0.114 for fMRIPrep (*P* = 0.562). **c,** The fMRI-to-anatomy (F2A) registration Dice coefficient was 0.952 ± 0.010 for NeuroImgAgent and 0.935 ± 0.016 for fMRIPrep (*P* < 0.001). The anatomy-to-template (A2T) registration Dice coefficient was 0.989 ± 0.001 for NeuroImgAgent and 0.983 ± 0.001 for fMRIPrep (*P* < 0.001). **d–f,** Comparison of preprocessing metrics between NeuroImgAgent and AFNI on rhesus macaque data (n = 30 pairs). d, Agreement between NeuroImgAgent and AFNI in mean FD and tSNR. e, QC–FC correlations. The mean absolute correlation coefficient (|r|) was 0.182 for NeuroImgAgent and 0.133 for AFNI (*P* = 0.782). f, The F2A registration Dice coefficient was 0.823 ± 0.080 for NeuroImgAgent and 0.641 ± 0.203 for AFNI (*P* < 0.001). The A2T registration Dice coefficient was 0.982 ± 0.004 for NeuroImgAgent and 0.972 ± 0.021 for AFNI (*P* = 0.020; n = 27 pairs for both comparisons). **g–i**, Comparison of preprocessing metrics between NeuroImgAgent and RABIES on rat data (n = 30 pairs). **g,** Agreement between NeuroImgAgent and RABIES in mean FD and tSNR. h, QC–FC correlations. The mean absolute correlation coefficient (|r|) was 0.179 for NeuroImgAgent and 0.172 for RABIES (*P* = 0.782). i, The F2A registration Dice coefficient was 0.888 ± 0.016 for NeuroImgAgent and 0.889 ± 0.019 for RABIES (*P* = 0.847). The A2T registration Dice coefficient was 0.940 ± 0.013 for NeuroImgAgent and 0.930 ± 0.009 for RABIES (*P* < 0.001). **j–l**, Comparison of preprocessing metrics between NeuroImgAgent and RABIES on mouse data. j, Agreement between NeuroImgAgent and RABIES in mean FD and tSNR across 38 observations. k, QC–FC correlations across 38 observations. The mean absolute correlation coefficient (|r|) was 0.316 for NeuroImgAgent and 0.294 for RABIES (*P* = 0.704). l, The F2A registration Dice coefficient was 0.932 ± 0.019 for NeuroImgAgent and 0.848 ± 0.075 for RABIES (*P* < 0.001). The A2T registration Dice coefficient was 0.938 ± 0.022 for NeuroImgAgent and 0.939 ± 0.027 for RABIES (*P* = 0.906; n = 23 pairs for both comparisons). QC–FC *P* values in b, e, h and k are from two-sided paired permutation tests and are Holm-adjusted across the four species. Dice *P* values in c, f, i and l are from two-sided paired t-tests. *P* < 0.05 (*) and *P* < 0.001 (***); n.s., not significant. Baseline denotes the reference workflow for each species.

We also examined the association between residual head motion and functional connectivity (QC–FC) as an exploratory, descriptive benchmark rather than a primary quality endpoint. Across all four species, mean absolute edgewise QC–FC correlations did not differ significantly between NeuroImgAgent and the corresponding reference workflow (two-sided paired permutation tests with Holm correction across the four species; all adjusted *P* ≥ 0.56; humans, n = 170; macaques, n = 30; rats, n = 30; mice, n = 38; **Fig. 2b,e,h,k)**. The comparison was also non-significant before correction in the resting-state-only human subset (n = 160). Because aggressive motion regression can lower QC–FC while also attenuating neural signal and reducing degrees of freedom, lower absolute QC–FC alone was not interpreted as evidence of better preprocessing.

Registration-related mask overlap was assessed in the subset with anatomical images using fMRI-to-anatomy (F2A) and anatomy-to-template (A2T) Dice coefficients (mean ± s.d.). In humans, F2A Dice was 0.952 ± 0.010 for NeuroImgAgent versus 0.935 ± 0.016 for fMRIPrep, and A2T Dice was 0.989 ± 0.001 versus 0.983 ± 0.001. Mean Dice was higher for NeuroImgAgent on both endpoints, with absolute differences of 0.017 and 0.006 (**Fig. 2c**; n = 170; both *P* < 0.001). In macaques, F2A Dice was 0.823 ± 0.080 for NeuroImgAgent and 0.641 ± 0.203 for AFNI, while A2T Dice was 0.982 ± 0.004 versus 0.972 ± 0.021 (**Fig. 2f**; n = 27 pairs; *P* < 0.001 and *P* = 0.020, respectively).

In rats, F2A Dice was 0.888 ± 0.016 for NeuroImgAgent and 0.889 ± 0.019 for RABIES, whereas A2T Dice was 0.940 ± 0.013 versus 0.930 ± 0.009 (**Fig. 2i**; n = 30; *P* = 0.847 for F2A and *P* < 0.001 for A2T). In mice, F2A Dice was 0.932 ± 0.019 for NeuroImgAgent versus 0.848 ± 0.075 for RABIES, and A2T Dice was 0.938 ± 0.022 versus 0.939 ± 0.027, with higher Dice for NeuroImgAgent for F2A and no significant difference for A2T (**Fig. 2l**; n = 23 pairs; *P* < 0.001 for F2A and *P* = 0.906 for A2T). Mean Dice was numerically higher for NeuroImgAgent in six of the eight comparisons. No statistically significant difference was detected for rat F2A or mouse A2T. These measurements characterize brain-mask overlap and do not directly assess internal anatomical alignment.

When the reference workflows were run on datasets as distributed, some cases were processed successfully by the reference workflows without manual preparation. Other cases showed incorrect image orientation or registration failure, or could not be processed because anatomical images were unavailable (**Extended Data Fig. 3**). These observations illustrate the input-handling challenges that NeuroImgAgent is designed to address through automated profiling, orientation correction and fallback mechanisms.

### VLM-guided quality control detects orientation, registration and brain-mask errors

We evaluated three VLMs for QC of image orientation, registration and brain masks: GPT-5.5 (OpenAI), Qwen3.8-Max (Alibaba) and the open-weight Llama 4 Scout (Meta). MRI orientation inspection and correction followed a gated workflow combining VLM assessment with geometric and template-based checks (**Fig. 3a**). After right–anterior–superior (RAS) standardization, the VLM assessed the anatomical axes and directions of each image. Cases that passed these checks proceeded directly to output. Flagged orientation errors triggered axis permutations and sign flips, whereas missing or low-confidence judgements triggered a geometry-based fallback. Figure 3b illustrates representative misoriented inputs and their outputs after VLM-guided correction for human and mouse data.

**Fig. 3:**
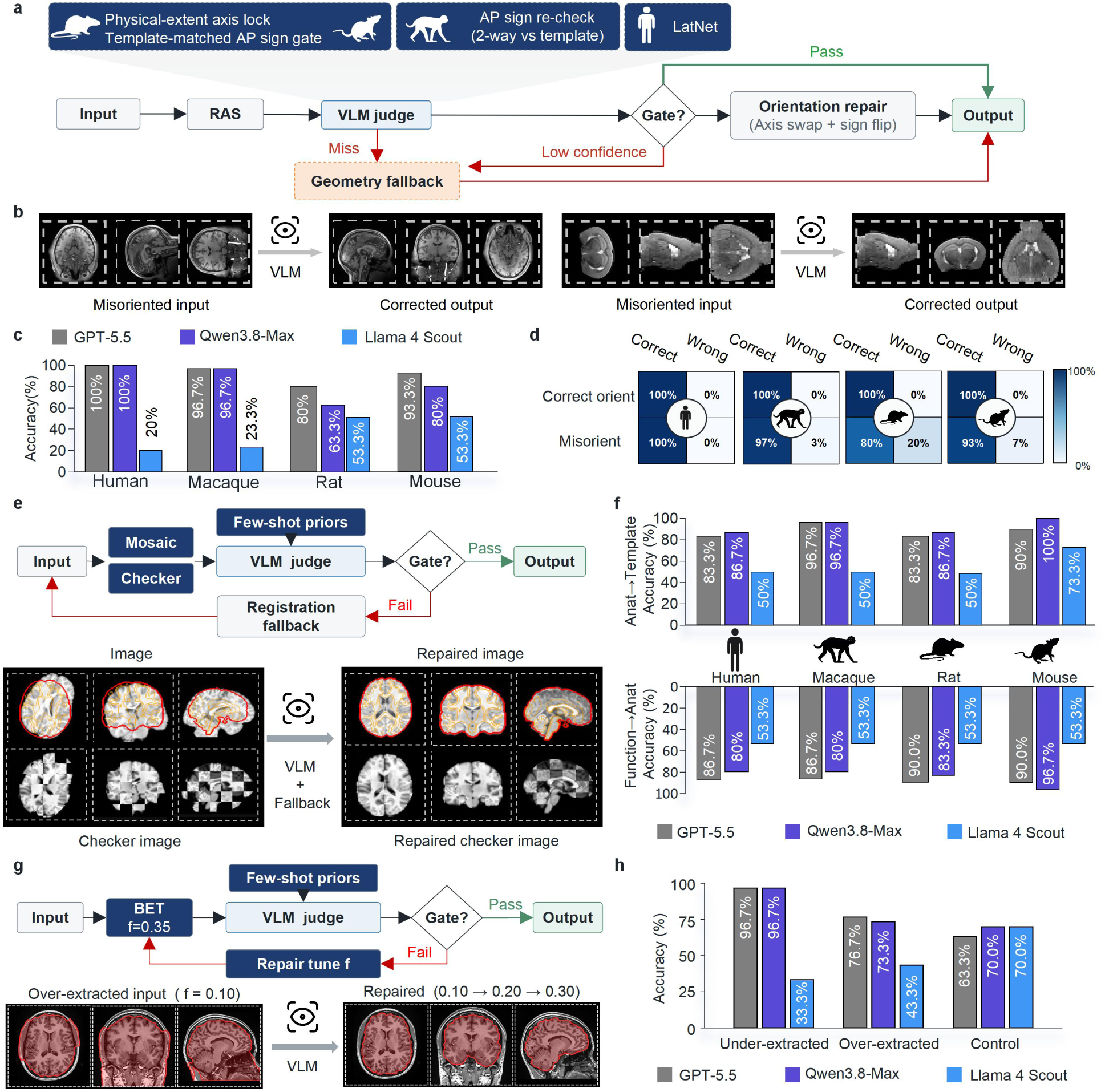
Vision–language model checkpoints for orientation, registration and brain-mask quality control. **a**, MRI orientation VLM inspection and correction workflow in NeuroImgAgent: after right–anterior–superior (RAS) standardization of the data, a vision–language model (VLM) assessed the anatomical axes and directions, supported by geometric checks and template-based checks. Axis permutations and sign flips were applied to correct orientation errors; missing or low-confidence judgements triggered a geometry-based fallback. **b**, Representative misoriented inputs and their corrected outputs are shown for human and mouse data. **c**, Exact orientation-correction accuracies of GPT-5.5, Qwen3.8-Max and Llama 4 Scout across 30 synthetically misoriented cases per species. For human data, the respective accuracies were 100%, 100% and 20%; for macaque data, 96.7%, 96.7% and 23.3%; for rat data, 80%, 63.3% and 53.3%; and for mouse data, 93.3%, 80% and 53.3%. **d**, Final orientation outcome of the GPT-5.5 workflow in humans, macaques, rats and mice. Rows indicate the ground-truth input class: correctly oriented controls (30 human, 30 macaque, 15 rat and 15 mouse cases) and misoriented inputs (30 cases per species). Columns indicate whether the delivered orientation matched the reference exactly (Correct output) or not (Wrong output). Values and colours give the percentage of each row (colour scale, 0–100%). **e**, Registration quality-control and repair workflow in NeuroImgAgent. A VLM with few-shot priors jointly evaluated the registration of functional images, structural images and template images using wireframe overlays and checkerboard images. A PASS judgement from the VLM indicated success, whereas a FAIL judgement triggered a registration fallback for re-registration, followed by reassessment. A representative failed human registration and its repaired result are shown. **f**, Accuracy of VLM-based registration quality assessment across four species. The upper and lower plots show anatomy-to-template (A2T) and fMRI-to-anatomy (F2A) registration assessments, respectively. Each species-specific dataset comprised 30 cases. For A2T registration, the accuracies of GPT-5.5, Qwen3.8-Max and Llama 4 Scout were, respectively, 83.3%, 86.7% and 50% for human; 96.7%, 96.7% and 50% for macaque; 83.3%, 86.7% and 50% for rat; and 90%, 100% and 73.3% for mouse. For F2A registration, the corresponding accuracies were 86.7%, 80% and 53.3% for human; 86.7%, 80% and 53.3% for macaque; 90%, 83.3% and 53.3% for rat; and 90%, 96.7% and 53.3% for mouse. **g**, Tiered quality control for human brain extraction. After checks for execution failure or empty masks, mask volume was evaluated against a range of 800–2,200 cm³, followed by VLM inspection. Over- or under-extraction prompted adjustment of the BET threshold f by 0.1 between runs, with up to three BET attempts. Representative repairs are shown. **h**, Quality-control classification accuracy for experimentally induced under-extraction, over-extraction and control masks without induced errors, with 30 cases per condition. Grey, purple and blue denote GPT-5.5, Qwen3.8-Max and Llama 4 Scout accuracy, respectively.

In the induced-error benchmark, exact correction was defined by recovery of the correct axis permutation and anterior–posterior and superior–inferior signs. With GPT-5.5, the workflow exactly corrected 30/30 human, 29/30 macaque, 24/30 rat and 28/30 mouse perturbations. The corresponding counts were 30/30, 29/30, 19/30 and 24/30 with Qwen3.8-Max, and 6/30, 7/30, 16/30 and 16/30 with Llama 4 Scout (**Fig. 3c**). On the 90 correctly oriented controls, the GPT-5.5 workflow delivered correctly oriented outputs in all cases (**Fig. 3d**). For PASS/REPAIR decisions on these controls, Qwen3.8-Max returned PASS for 89 of 90, whereas Llama 4 Scout incorrectly flagged 48 of 90 for repair, corresponding to a 53.3% false-alarm rate (**Extended Data Fig. 4a**).

For registration QC, a VLM assessed wireframe overlays and checkerboard views using same-species PASS/FAIL examples as few-shot references. A FAIL judgement triggered a registration fallback followed by reassessment (**Fig. 3e**). Registration classification was evaluated using a dedicated 120-case test set for each of F2A and A2T. Each set included 30 cases per species, comprising 15 true PASS and 15 true FAIL cases. For F2A registration, GPT-5.5, Qwen3.8-Max and Llama 4 Scout correctly classified 106/120 (88.3%), 102/120 (85.0%) and 64/120 (53.3%) cases, respectively. For A2T registration, the corresponding counts were 106/120 (88.3%), 111/120 (92.5%) and 67/120 (55.8%) (**Fig. 3f**). Error profiles also differed between models. For F2A, GPT-5.5 falsely rejected 6 correctly aligned registrations, compared with 11 for Qwen3.8-Max. For A2T, Qwen3.8-Max falsely accepted 9 of 60 failed registrations, compared with 14 of 60 for GPT-5.5 (**Extended Data Fig. 4c**).

Brain-mask classification and tiered repair were evaluated as separate experiments using the same 30 human subjects under control, induced under-extraction and induced over-extraction conditions. In the full repair workflow, a deterministic volume gate could intercept a mask before VLM review, and failed QC checks could trigger parameter adjustment in FSL’s Brain Extraction Tool (BET) or a configured fallback (**Fig. 3g**). For tiered QC of the initial masks before repair, the numbers correctly classified as control, under-extraction and over-extraction were 19/30, 29/30 and 23/30 with GPT-5.5. The corresponding counts were 21/30, 29/30 and 22/30 with Qwen3.8-Max, and 21/30, 10/30 and 13/30 with Llama 4 Scout (**Fig. 3h**). The volume gate intercepted the same 13/30 over-extracted masks in all three configurations. For Llama 4 Scout, the volume gate accounted for all 13 correct over-extraction classifications, whereas the separate VLM-only experiment yielded 0/30 correct over-extraction classifications (**Extended Data Fig. 4d and Supplementary Table 3**).

### Profiling and quality control improve species identification and output usability

Replacing the specialized profiler with a generic alternative exposed substantial species-identification errors. With the profiler in place (Full), all 120 cases (30 per species) were assigned the correct species. In the −Profiler configuration, which used a generic evidence collector followed by a single GPT-5.5 call, 68 of 120 cases (56.7%) were correctly identified, whereas 24 (20.0%) were misclassified and 28 (23.3%) received no species decision (**Fig. 4a**). All misclassifications involved rat cases labelled as mice, leaving only one of 30 rat cases correctly identified, compared with 25 human, 17 macaque and 25 mouse cases (**Fig. 4b**). This concentration of misclassifications highlights rat–mouse discrimination as a particular vulnerability of the −Profiler configuration.

**Fig. 4:**
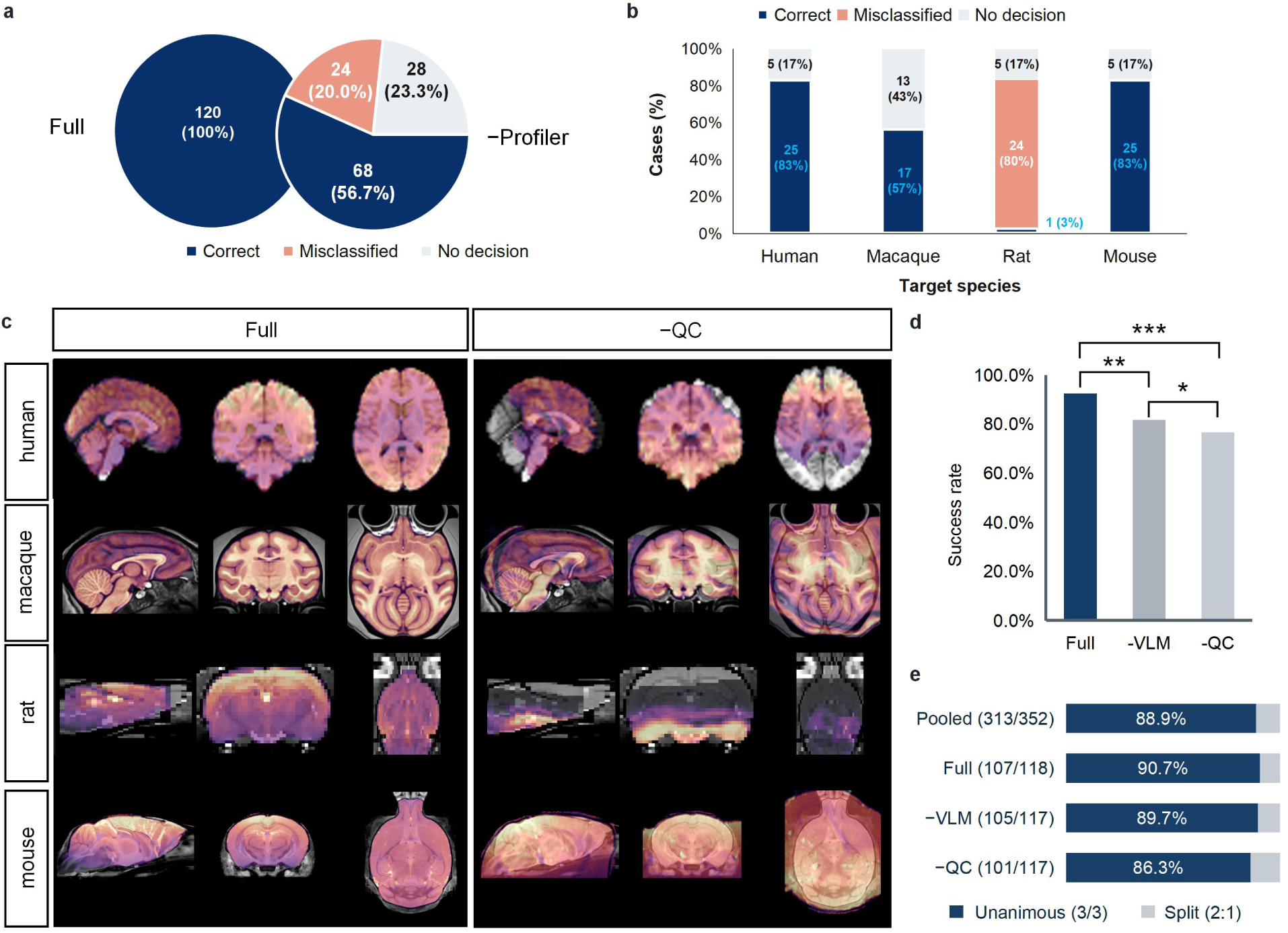
Ablation of profiling and quality control in NeuroImgAgent. **a**, Species identification in 120 cases (30 per species) with and without the specialized profiler. With the profiler (Full; left), all 120 cases (100%) were assigned the correct species. When the profiler was replaced by a generic evidence collector followed by a single GPT-5.5 call (−Profiler; right), 68 cases (56.7%) were correctly identified, 24 (20.0%) were misclassified and 28 (23.3%) yielded no species decision. **b**, Outcome distributions in the −Profiler configuration, stratified by target species (n = 30 per species). Correct identification was achieved for 25 human, 17 macaque, 1 rat and 25 mouse cases. All 24 misclassifications involved rat cases classified as mouse. Five human, 13 macaque, 5 rat and 5 mouse cases yielded no decision. Numbers within segments indicate case counts and percentages. “No decision” denotes that the model did not assign a species label. **c**, Representative sagittal, coronal and axial registration overlays for human, macaque, rat and mouse challenge cases, comparing Full with the indicated ablated configuration. Registered images are shown in colour over greyscale references. **d**, Overall success rates for Full, −VLM and −QC were 92.5% (111/120), 81.7% (98/120) and 76.7% (92/120), respectively, across 120 paired challenge cases comprising 30 cases per species. Success was defined by majority expert assessment of overall usability; runs without evaluable outputs were counted as unsuccessful. Comparisons used two-sided exact McNemar tests with Holm correction across the three pairwise comparisons. *P* < 0.05 (*), *P* < 0.01 (**) and *P* < 0.001 (***), based on adjusted *P* values. **e**, Agreement among three expert reviewers on overall usability, counted by rated case–configuration entries. Dark blue indicates unanimous judgements (3/3), including unanimous PASS and FAIL, and light grey indicates split judgements (2:1).

We next examined QC ablations in 120 paired challenge cases, with 30 cases per species. Output usability was determined by the majority judgement of three expert reviewers blinded to configuration. Runs without evaluable outputs were counted as unsuccessful. Full achieved an expert-assessed usability rate of 92.5%, compared with 81.7% for −VLM and 76.7% for −QC (**Fig. 4d**). Full exceeded both ablated configurations (Holm-adjusted *P* = 0.002 and *P* < 0.001, respectively), and −VLM exceeded −QC (adjusted *P* = 0.031; two-sided exact McNemar tests). Representative overlays illustrated poorer anatomical alignment after QC removal (**Fig. 4c**). Judgements from the three reviewers were unanimous for 313 of 352 rated case-configuration entries (88.9%; Fig. 4e), with shared image plates and their ratings counted once in each represented configuration. Together, these results support a role for profiling in species identification and an association between the full configuration and higher output usability.

Further analysis showed that Full also limited the delivery of unusable outputs without warning. Such deliveries occurred in 5/120 cases (4.2%) with Full, compared with 15/120 (12.5%) with −VLM and 22/120 (18.3%) with −QC (Holm-adjusted *P* = 0.013 and *P* < 0.001, respectively; **Extended Data Fig. 5a,b**). In the 24 orientation-challenge cases, Full achieved usability in 20 cases (83.3%), compared with 10 (41.7%) for −VLM and 6 (25.0%) for −QC (**Extended Data Fig. 5c,d**). Overall, the full configuration was associated with higher output usability and fewer unusable outputs delivered without warning. This usability advantage was also observed in the orientation-challenge subset.

### NeuroAgentBench evaluates agent capabilities and workflow execution

NeuroAgentBench included 28 L1, 24 L2, 9 L3 and 14 L4 cases per model, with failures and dependency-blocked cases retained in the pass-rate denominators (**Fig. 5a,b and Extended Data Fig. 6**). With GPT-5.5, pass rates for L1-L4 were 100.0%, 100.0%, 88.9% and 78.6%, respectively. The corresponding rates were 100.0%, 100.0%, 66.7% and 28.6% with Qwen3.8-Max, and 96.4%, 95.8%, 77.8% and 35.7% with Llama 4 Scout. Within this benchmark, high pass rates for intent recognition and planning did not consistently extend to fault recovery and tool construction.

**Fig. 5:**
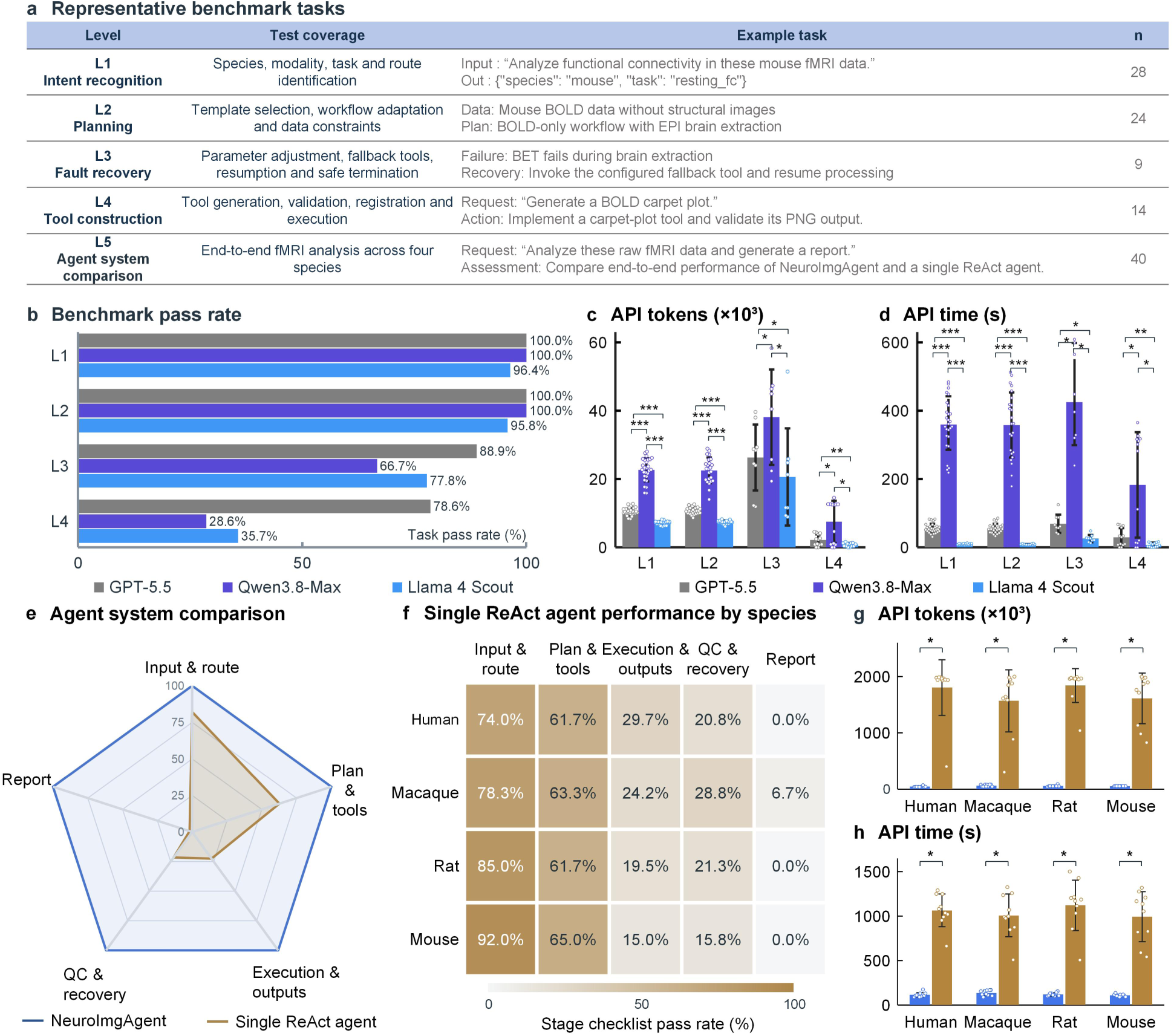
NeuroAgentBench: task capabilities and end-to-end performance. **a**, The benchmark of NeuroImgAgent, termed NeuroAgentBench, comprises representative tasks spanning intent recognition (L1), planning (L2), fault recovery (L3), tool construction (L4) and agent-system evaluation (L5). **b**, Case-level pass rates for GPT-5.5, Qwen3.8-Max and Llama 4 Scout (grey, purple and blue, respectively). Failures and dependency-blocked cases remain in the denominators. **c,d**, Per-case API token consumption (input + output, ×10³; c) and cumulative API response time (s; d). Bars, error bars and points indicate means, sample standard deviations and individual cases, respectively. Within each level, group n follows GPT-5.5/Qwen3.8-Max/Llama 4 Scout. The corresponding paired n follows GPT-5.5–Qwen3.8-Max/GPT-5.5–Llama 4 Scout/Qwen3.8-Max–Llama 4 Scout. Group n in c was 28/27/28 (L1), 24/24/24 (L2), 9/8/8 (L3) and 14/11/14 (L4). Paired n in c was 27/28/27 (L1), 24/24/24 (L2), 8/8/8 (L3) and 11/14/11 (L4). Group n in d was 28/27/28 (L1), 24/24/24 (L2), 9/8/9 (L3) and 14/11/14 (L4). Paired n in d was 27/28/27 (L1), 24/24/24 (L2), 8/9/8 (L3) and 11/14/11 (L4). Holm-adjusted *P* values in c, reported in the comparison order defined above, were all *P* < 0.001 at L1 and L2; *P* = 0.039, *P* = 0.039 and *P* = 0.039 at L3; and *P* = 0.018, *P* = 0.010 and *P* = 0.018 at L4. Holm-adjusted *P* values in d were all *P* < 0.001 at L1 and L2; *P* = 0.039, *P* = 0.027 and *P* = 0.039 at L3; and *P* = 0.018, *P* = 0.010 and *P* = 0.018 at L4. **e**, Comparison of NeuroImgAgent and a single ReAct agent across input and routing, planning and tools, execution and outputs, quality control (QC) and recovery, and reporting. The single-agent radar profile represents equally weighted species means. **f**, Species-specific stage checklist scores for the GPT-5.5-based single ReAct agent across 40 fMRI cases, with 10 cases each from humans, macaques, rats and mice. Checkpoint scores are fractions of satisfied checklist items, averaged equally within stages and across cases. **g,h**, L5 main-agent input-plus-output tokens (g) and cumulative API response time (h), using GPT-5.5. Bars show means ± sample s.d., and points show all selected cases (n = 10 per species and system). For both g and h, the comparisons between NeuroImgAgent and the single ReAct agent yielded Holm-adjusted *P* = 0.027 for human, macaque, rat and mouse. Tool-internal VLM calls are excluded. Statistical comparisons used two-sided exact Wilcoxon signed-rank tests paired by case or subject. All 32 comparisons were jointly adjusted using Holm’s method. Asterisks were assigned using the unrounded adjusted values: \**P* < 0.05, \*\**P* < 0.01 and \*\*\**P* < 0.001.

For L1-L4, API token consumption and cumulative remote-model response time were summarized for cases with complete usage records (**Fig. 5c,d**). Failed cases with complete records and verified zero-call executions were retained. Dependency-blocked cases and cases with incomplete usage records were excluded. API time excluded local computation and failed-request waiting. Level-specific sample sizes are reported in the figure legend, and the thresholds applied to resource summaries are described in the Methods. Failures and dependency-blocked cases remained in the denominators for the pass-rate analysis.

The Toolsmith workflow specifies inputs, generates and validates candidate code, and registers validated tools for execution (**Extended Data Fig. 7a,b**). In a separate offline illustration of QC summarization, an archived standardized DVARS trace from human subject sub-70080 was used to group 30 flagged frames into nine continuous outlier periods, corresponding to a total duration of 60 s (**Extended Data Fig. 7c**).

We compared NeuroImgAgent with a single ReAct agent using GPT-5.5 on 40 matched cases, with 10 cases per species (**Fig. 5e,f**). Both systems were evaluated using the same stage-specific checklist covering input and routing, planning and tool selection, execution and outputs, QC and recovery, and reporting. NeuroImgAgent achieved stage scores of 100%, whereas the single agent achieved 74.0%, 61.7%, 29.7%, 20.8% and 0.0% for humans; 78.3%, 63.3%, 24.2%, 28.8% and 6.7% for macaques; 85.0%, 61.7%, 19.5%, 21.3% and 0.0% for rats; and 92.0%, 65.0%, 15.0%, 15.8% and 0.0% for mice, respectively. The corresponding comparisons of selected main-agent token use and API response time are shown in **Fig. 5g,h**. A descriptive analysis of 40 single-agent development attempts further characterized failure modes, recovery checks and final deliverables (**Extended Data Fig. 8**).

### Midnight Scan Club reanalysis recovers established patterns of network similarity

Beyond preprocessing metrics, we reanalysed the Midnight Scan Club dataset to assess whether NeuroImgAgent reproduced previously reported patterns of functional-network similarity. The analysis included nine participants and yielded 90 functional-connectivity matrices spanning five states and two session halves per participant. Consistent with prior work^22,23^, participant-level comparisons showed higher functional-network similarity in the Individual than in the Group condition, and in the Individual-by-Task than in the Individual condition (both q < 0.001). Normalized relative contributions to functional-network similarity were largest for the Group (40.2%) and Individual (33.1%) components, followed by the Individual-by-Task (18.1%), Individual-by-Session (4.9%) and Task (3.6%) components (**Extended Data Fig. 9**). These findings show that NeuroImgAgent reproduced previously reported individual and individual-by-task patterns of functional-network similarity within the Midnight Scan Club dataset.

## Discussion

NeuroImgAgent provides a shared framework for fMRI analysis across humans, macaques, rats and mice by coordinating species-aware workflow selection, execution and recovery through reviewable evidence. The framework retains species-specific templates, parameters and tools while building on established preprocessing pipelines^2–4^. Across 268 paired cases, FD and tSNR estimates were broadly similar to those of the reference workflows. Among cases with anatomical images, mean brain-mask overlap after registration was numerically higher for NeuroImgAgent in six of the eight species-by-endpoint comparisons (**Fig. 2**). The profiler ablation further showed that replacing the specialized profiler with a generic evidence collector followed by a single GPT-5.5 call introduced substantial species-identification errors, with all misclassifications involving rats labelled as mice (**Fig. 4a,b**). This finding supports the value of specialized profiling for selecting species-appropriate workflows.

Evidence-gated QC links the assessment of intermediate outputs to decisions about recovery and downstream execution. In the challenge cohort, the full configuration improved expert-rated output usability and reduced the delivery of unusable outputs without warning (**Fig. 4d and Extended Data Fig. 5a,b**). Quantitative checks and visual assessment guide predefined repairs and reassessment, while downstream execution stops when detected fatal QC failures remain unresolved. Recorded parameters, tool versions and recovery histories allow researchers to review how outputs were produced and how detected problems were handled.

Related scientific agents illustrate complementary approaches to structuring and checking automated analyses. CellAgent assigns specialized planning, execution and evaluation roles for single-cell analysis^16^, and GeneAgent verifies gene-set interpretations against curated biological databases^18^. CopilotJ connects conversational multi-agent assistance with ImageJ and supports the validation and reuse of analysis workflows^20^. NeuroImgAgent focuses on species-aware workflow selection and evidence-gated assessment of image orientation, brain masks and registration in cross-species fMRI. Its natural-language interface and human approval of proposed workflows are intended to make analysis accessible to researchers with limited programming experience.

NeuroAgentBench showed that strong intent recognition and planning did not consistently translate into reliable recovery or tool construction. The Midnight Scan Club reanalysis provided a complementary downstream test by reproducing previously reported individual and individual-by-task patterns of functional-network similarity. Together, these evaluations assess both operational reliability and the reproduction of an established neuroimaging analysis.

Several limitations define the scope of these findings. Visual judgements varied across models and species, with false acceptances of failed registrations and false alarms on control brain masks. Some unusable outputs were still delivered without warning under the full configuration. The brain-mask assessment and L3 recovery benchmark used only human data. Controlled perturbations enable targeted testing, but performance on naturally occurring failures requires further evaluation. Dice measures mask overlap rather than internal anatomical alignment, and non-significant QC–FC differences do not establish equivalence between workflows. The Midnight Scan Club reanalysis used the original dataset and does not establish generalizability to independent cohorts or biological validity across species. The effects of NeuroImgAgent on user time, workload and accessibility also remain to be measured directly.

Looking ahead, we plan to extend NeuroImgAgent beyond fMRI to other MRI modalities, including diffusion MRI and magnetic resonance spectroscopy. The separation of profiling, planning, evidence-gated QC and reporting from the registered tools provides a basis for these extensions. Each extension will require modality-specific tools, QC criteria and benchmark tasks, together with comparisons against established workflows and validation using relevant scientific endpoints. The present results establish a framework for cross-species fMRI analysis with traceable processing decisions and improved output usability in the evaluated challenge cases.

## Methods

### Datasets, software and ethics

We assembled public and institutional MRI data spanning humans, macaques, rats and mice (**Supplementary Table 1**). NeuroImgAgent processed 175 human subjects; 170 were included in the paired fMRIPrep comparison because five ds000030 subjects lacked a T1-weighted image and could not be handled by fMRIPrep. The paired cohort comprised 117 UCLA Consortium for Neuropsychiatric Phenomics cases (OpenNeuro ds000030), 15 HNU CoRR cases, 10 Human Connectome Project 3 T REST1 cases, 10 Midnight Scan Club cases (ds000224, version 1.0.4), 10 movie-viewing cases (ds000228, local snapshot version 1.1.0) and 8 UMR-5T cases. The macaque dataset comprised 30 cases. Macaque data comprised 7 AFNI demonstration cases, 10 Oxford cases, 10 University of Wisconsin–Madison cases and 3 NeuroSpin BOLD-only cases. The rat data consisted of 30 cases. Rat data comprised six cases from each of five sites in OpenNeuro ds004116. The mouse cohort comprised 38 cases. Mouse data comprised 5 cases each from ds001653, ds004125 and ds001890, 15 BOLD-only cases from ds001720 and 8 cases from an institutional awake-mouse cohort.

Registration used MNI152NLin2009cAsym, NMT, Fischer 344 and DSURQE reference spaces for humans, macaques, rats and mice, respectively. NeuroImgAgent orchestrated versioned tools from ANTs, FSL, AFNI and SimpleITK, together with NiBabel, Nilearn, NumPy, SciPy and related Python libraries (**Supplementary Table 5**). Reference comparisons used fMRIPrep 24.1.1, AFNI 26.1.00 and RABIES 0.5.1. Dataset access conditions, acquisition summaries and template licences are listed in Supplementary Table 1; reference-workflow commands are listed in **Supplementary Table 2**.

Public datasets were analysed in accordance with their source-specific access terms and the ethical approvals under which they were collected. The UMR-5T study was approved by the Medical Ethics Committee of the Shenzhen Institutes of Advanced Technology, Chinese Academy of Sciences (SIAT-IRB-260715-H1094). All participants provided written informed consent in accordance with the Declaration of Helsinki. The institutional awake-mouse study was approved by the Institutional Animal Care and Use Committee of the Shenzhen Institutes of Advanced Technology, Chinese Academy of Sciences (SIAT-IACUC-240702-YGS-LY-A2838).

### Language models and inference settings

We evaluated GPT-5.5, Qwen3.8-Max and Llama 4 Scout through OpenAI-compatible interfaces. Non-GPT branches requested temperature=0; GPT-5.5 requests omitted temperature and specified reasoning_effort=medium. Generation budgets were set with max_completion_tokens: 16,384 tokens for routine analysis and 2,048 tokens for registration. Qwen3.8-Max responses reported reasoning tokens, although no reasoning controls were requested. Temperature 0 and greedy decoding do not guarantee identical outputs across repeated calls.

### Data ingestion, profiling and species identification

NeuroImgAgent supports multiple neuroimaging data formats, including DICOM and NIfTI. For non-BIDS inputs, the ingestion layer indexes the available imaging files and associated metadata sidecars and converts them into a normalized internal representation without modifying the source data. BIDS-compatible inputs follow the standard dataset and entity conventions^1^.

For each dataset, the profiler inventories available structural images, functional images, field maps and event files, and records the image dimensions, voxel sizes, orientation, number of time points and relevant metadata. It then determines whether the modalities needed for the requested analysis are present and generates a structured dataset profile that distinguishes available, missing and unresolved inputs. Species identification integrates declared metadata with acquisition geometry, estimated brain size and image-derived evidence.

### Workflow planning and user approval

The planner converted each natural-language request into a structured specification containing the analysis task, imaging modalities, requested outputs and user-defined constraints. It selected a compatible pipeline template, instantiated the permitted species-dependent options, verified the availability of the required tools and constructed a directed acyclic graph after validating the declared input–output dependencies of each node and the coverage of required outputs. Before execution, the proposed graph, selected operations and expected outputs were presented for review by a human. Execution proceeded only after approval. Requested modifications, the approved graph and the final resolved configuration were retained in the run record. The experiments reported here used the approved configurations associated with the corresponding source figure packages.

### Execution, checkpoints and provenance

The executor parallelized work across subjects within the configured resource limit. Within a subject, a node ran only after its declared upstream dependencies completed successfully. For each node, the system recorded input and output identities, resolved parameters, tool version, execution status, timestamps and QC outcome.

Successful nodes generated checkpoints, allowing interrupted runs and recovery attempts to resume from the affected branch without repeating unrelated completed work. A circuit breaker monitored repeated occurrences of the same failure across subjects and, once a configured threshold was reached, prevented new cases from launching. Run-level reports linked final products to the corresponding execution and QC evidence.

### Quality control, recovery and reporting

QC was implemented as an explicit gate between processing nodes. Each tool manifest declared required outputs, quantitative metrics, artifact checks, decision thresholds and rule severity. File-integrity checks, numerical measurements and standardized image mosaics were assembled into an evidence bundle. PASS and non-fatal WARN outcomes permitted continued execution; a fatal FAIL closed the downstream gate. Recovery was restricted to predefined parameter adjustments, upstream retuning or registered fallback procedures. The affected node was then re-executed and reassessed, with retry budgets and circuit-breaker thresholds limiting the loop.

VLM review was combined with task-specific deterministic evidence for image orientation, registration and brain masking. Orientation review compared candidate slice mosaics, with geometric and template-profile fallbacks when the model call failed, could not be parsed or did not meet decision requirements. Registration QC evaluated standardized overlays and checkerboards after catastrophic objective failures had been rejected. Brain-mask QC screened execution status, empty masks and prespecified volume bounds before VLM assessment of over-or under-extraction. All model responses, evidence images, deterministic decisions, fallback selections and retry histories were retained. A VLM verdict, a workflow-log PASS and a fallback handoff were stored as separate events and were not treated as interchangeable validation outcomes.

### Preparation of inputs for reference workflows

Manual preparation for the reference workflows comprised reorientation of rat, anatomical-mouse and awake-mouse images to a standard axis convention, organization of awake-mouse scanner files into BIDS, use of an EPI-derived anatomical reference for macaques without T1-weighted images, and conversion of the HNU and UMR-5T human data to BIDS (file renaming for HNU; DICOM-to-NIfTI conversion for UMR-5T); the remaining cohorts were processed as distributed.

### Human data and comparison with fMRIPrep

The human benchmark comprised 170 paired comparison units from six datasets: the UCLA Consortium for Neuropsychiatric Phenomics dataset (OpenNeuro ds000030)^24^, HNU1 from the Consortium for Reliability and Reproducibility, Human Connectome Project 3 T REST1, the Midnight Scan Club (OpenNeuro ds000224)^22^, the movie-viewing dataset OpenNeuro ds000228^25^ and the in-house UMR-5T dataset. Each case was processed independently by NeuroImgAgent and fMRIPrep 24.1.1, and volumetric derivatives were evaluated in MNI152NLin2009cAsym space. FD and tSNR were calculated for all 170 paired units. QC–FC analyses included 170 paired units; for each edge, the Pearson correlation between subject-level mean FD and edge functional connectivity was computed across eligible records, and the endpoint was the mean absolute edgewise correlation, tested with two-sided paired permutation tests (9,999 permutations) that exchanged whole records. For fMRI-to-anatomy (F2A) alignment, each workflow’s EPI brain mask in MNI space was compared with that same workflow’s transformed anatomical mask; for anatomy-to-template (A2T) alignment, each workflow’s template-space anatomical mask was compared with the same fixed template brain mask. Dice values are reported as mean ± s.d. across the 170 paired cases and tested with two-sided paired t-tests.

### Macaque data and comparison with AFNI

The macaque benchmark included data from the Oxford, University of Wisconsin–Madison and NeuroSpin cohorts, together with the AFNI macaque demonstration dataset; the data were obtained through PRIME-DE or the corresponding public AFNI resources^26–31^. The reference workflow reproduced the AFNI MACAQUE_DEMO_REST workflow, comprising @animal_warper followed by afni_proc.py, with registration to the symmetric NMT v2.1 template at 0.5-mm resolution^4,32^.

FD and tSNR were evaluated for 30 paired runs. The primary pooled QC–FC analysis included all 30 runs (29 resting-state scans and one task scan; 231 edges). Registration analyses included 27 runs with T1-weighted images; three BOLD-only runs were ineligible for registration evaluation. One rest–task pair shared a T1-weighted image; the two runs were analysed as separate runs, so registration statistics and paired tests used all 27 runs. For F2A alignment, each workflow’s EPI mask on the paired AFNI evaluation grid was compared with that same workflow’s transformed anatomical mask; for A2T alignment, each workflow’s transformed anatomical mask was compared with the same fixed AFNI mask_group reference derived from the NMT v2.1 template. Dice values are reported as mean ± s.d. across the 27 runs.

### Rat data and comparison with RABIES

The rat comparison included 30 paired cases from OpenNeuro ds004116, with six cases from each of five sites operating at 4.7–17.15 T. NeuroImgAgent outputs were compared with precomputed outputs from RABIES 0.5.1 in Fischer 344 template space. FD, tSNR, QC–FC, F2A registration and A2T registration included all 30 pairs. Registration was evaluated with the Dice coefficient for two endpoints. For F2A alignment, each workflow’s anatomy-in-EPI mask was compared with its own EPI mask in the native EPI space. For A2T alignment, a common RS2Net native anatomical mask was fixed for each animal, transformed by each workflow’s saved transforms into the Fischer 344 template grid, and both transformed masks were compared with the same fixed template brain mask. Dice values are reported as mean ± s.d. across the 30 paired cases and tested with two-sided paired t-tests.

### Mouse data and comparison with RABIES

The mouse benchmark comprised 38 cases: 5 each from ds001653, ds004125 and ds001890 with anatomical images; 15 BOLD-only cases from ds001720; and 8 cases from an institutional awake-mouse cohort. NeuroImgAgent outputs were compared with precomputed outputs from RABIES 0.5.1. FD, tSNR and QC–FC included all 38 cases. F2A and A2T comparisons were evaluated in DSURQE space for the 23 cases with anatomical images; the 15 BOLD-only cases from ds001720 were excluded from registration evaluation because they lacked anatomical images.

### NeuroAgentBench

NeuroAgentBench defines five levels of increasing operational difficulty. L1 tests intent recognition and routing, L2 data-aware workflow planning, L3 diagnosis and bounded recovery from controlled failures, L4 construction, reuse, clarification or rejection of missing-tool requests, and L5 matched end-to-end comparison of single- and multi-agent systems (**Supplementary Table 4**). The completed evaluation reported here comprised 28 L1, 24 L2, 9 L3 and 14 L4 cases per model, together with an L5 end-to-end comparison that included 10 cases per species and system. A case passed only if all blocking, level-specific criteria were met; failures and dependency-blocked cases remained in the denominators. For L5 resource statistics, NeuroImgAgent retained the profiler, planner, first report writer and first report auditor of each case, whereas the single ReAct agent retained its sequential decision calls.

GPT-5.5, Qwen3.8-Max and Llama 4 Scout were evaluated. Token use was calculated from provider-reported input and output tokens across recorded calls, including retries. API time was calculated as the sum of remote model-call durations and excluded local computation and failed-request waiting. Records exceeding 30,000 tokens (L1, L2 and L4), 60,000 tokens (L3) or 600 s were excluded from the plotted statistics; a sensitivity analysis restoring records excluded only by these thresholds did not change the inferences. Resource summaries retained failures with complete records and verified zero-call runs but excluded incomplete usage records and blocked dependencies. The human single-agent token total for one case is a lower bound because two failed requests had unknown usage.

Group sizes were 28/27/28 (L1), 24/24/24 (L2), 9/8/8 (L3 tokens), 9/8/9 (L3 time) and 14/11/14 (L4) for GPT-5.5/Qwen3.8-Max/Llama-4-Scout; paired analyses used 27/28/27 (L1), 24/24/24 (L2), 8/8/8 (L3 tokens), 8/9/8 (L3 time) and 11/14/11 (L4) pairs. Statistical tests were two-sided exact Wilcoxon signed-rank tests paired by case or subject, with zero differences excluded and tied ranks averaged; all 32 comparisons were jointly adjusted using Holm’s method.

### Orientation correction experiment

We first standardized the source images to a verified RAS orientation. Synthetic orientation errors were then introduced by applying one of the five non-identity permutations of the three spatial axes, with additional AP or SI reversals. Each species contributed 30 misoriented cases. Correct-orientation controls included 30 human, 30 macaque, 15 rat and 15 mouse cases. For human images, the VLM first identified the LR axis from three candidate planes and then selected among eight AP/SI orientation hypotheses referenced to an MNI template, with LR polarity assessed using LatNet ^10^. Macaque images followed the same two-stage procedure using the NMT template, followed by a focused two-candidate AP-sign check. For rat and mouse images, an RS2 brain mask identified the AP, SI and LR axes from their physical extents; AP polarity was determined by comparison with Fischer344 or DSURQE template profiles. The VLM determined SI polarity directly for mouse images and selected between the current and SI-reversed candidates for rat images, with template-profile fallback when no usable VLM decision was returned. The predicted permutation and signs were then applied and verified against the registered inverse transformation. Exact correction accuracy was calculated as the number of cases with the correct axis permutation and AP/SI signs divided by 30; any error in these components counted the case as incorrect. Correct-orientation controls were excluded from the correction accuracy calculation. For the GPT-5.5 workflow, final orientation outcomes were summarized separately for correctly oriented controls and misoriented inputs, according to whether the delivered orientation matched the reference exactly (Correct) or not (Wrong) (Fig. 3d).

### Registration quality-control experiment

Registration-VLM test cases were represented by axial, coronal and sagittal overlays and a checkerboard image. The VLM received one same-species PASS/FAIL reference pair from a held-out subject, followed by the anonymized test images, and rated brain-boundary alignment, internal structures, signal dropout, systematic displacement and residual distortion on a 1–5 scale. Scores were averaged and rounded half up; an overall score of at least 3 was classified as PASS. F2A and A2T evaluations each used a dedicated 120-case test set: 30 cases per species, with 15 correctly aligned and 15 deterministically perturbed registrations. Accuracy and the numbers of false rejections and false acceptances were summarized descriptively.

### Brain extraction quality-control experiment

Human T1-weighted images were initially processed with FSL BET using a fractional intensity threshold of 0.35. Outputs were checked sequentially for execution failure, empty masks and brain-mask volumes outside 800–2,200 cm³; masks passing these checks underwent VLM classification for over- or under-extraction. The BET threshold was increased by 0.1 for over-extraction and decreased by 0.1 for under-extraction, followed by reprocessing. Up to three BET attempts were allowed before invoking the configured fallback. Classification performance was evaluated using 90 masks from the same 30 subjects—30 under-extracted, 30 over-extracted and 30 controls—for each model.

### Profiler and quality-control ablations

The profiler ablation evaluated species identification in 120 cases, comprising 30 human, 30 macaque, 30 rat and 30 mouse cases. In this configuration, the specialized profiler was replaced by a generic evidence collector followed by a single GPT-5.5 call (single-shot profiling). Outcomes were classified as correct species identification, misclassification or no decision, where no decision was defined as the absence of an assigned species label. Counts and proportions were summarized overall and by target species.

The QC ablation included 120 challenge cases, with 30 cases per species and paired results for the Full, −VLM and −QC configurations. Challenge cases were derived from 120 source subjects by introducing controlled perturbations into the inputs. Overall output usability was determined by the majority judgement of three reviewers with experience in fMRI preprocessing, who were blinded to configuration. A case was classified as successful when at least two reviewers judged its output usable. An output was rated usable when image orientation was correct, the brain mask showed no gross over- or under-extraction, and both registration steps showed no gross misalignment. Runs without evaluable outputs were classified as unsuccessful and retained in the denominator of 120 cases for each configuration.

Reviewer agreement was summarized as the proportion of rated case–configuration entries with unanimous judgements (3/3), including unanimous PASS and FAIL judgements; the remaining entries had split judgements (2:1). Agreement denominators were 118 for Full, 117 for −VLM and 117 for −QC. Pooled agreement was calculated by summing the configuration-specific numerators and denominators without cross-configuration deduplication, yielding 352 rated case–configuration entries. Shared plates and their associated ratings contributed once to each represented configuration. These entries were not treated as independent reviews.

Success rates were compared using two-sided exact McNemar tests, paired by case. Holm correction was applied to the three pairwise comparisons, with adjusted *P* < 0.05 considered statistically significant. Reviewer-agreement proportions were reported descriptively.

### Functional-network similarity decomposition

We analysed the Midnight Scan Club dataset (OpenNeuro ds000224, version 1.0.4), including nine participants (MSC01–MSC07, MSC09 and MSC10); MSC08 was excluded a priori because of high head motion and reported sleep. Preprocessed BOLD data were sampled onto participant-specific cortical surfaces, resampled to fsLR32k, smoothed with a Gaussian kernel (σ = 2.55 mm) and averaged within the 333-parcel Gordon atlas^33^. Parcel time series were generated for rest, motor, semantic, coherence and memory states and concatenated within two independent session halves (ordinal sessions 1–5 and 6–10), yielding 90 half-state functional-connectivity matrices (9 participants × 5 states × 2 halves). Functional connectivity (FC) was calculated as the Pearson correlation between parcel time series. Each matrix was represented by its Fisher-z-transformed upper-triangular edges; functional-network similarity was calculated as the Pearson correlation between edge vectors and then Fisher-transformed. Pairwise similarities were classified as Group, Task, Individual, Individual-by-Session or Individual-by-Task according to shared participant, state and data half^23^. Incremental contributions were calculated relative to the corresponding baseline, normalized within each participant and averaged across participants. The same decomposition was applied to each parcel’s connectivity profile and projected onto lateral and medial fsLR32k surfaces. Extended Data Fig. 9 shows the 90 × 90 similarity matrix, participant-level matched-factor effects shown as means ± 1 s.d., normalized effect composition and parcelwise cortical maps. Effects were compared using two-sided paired t-tests at the participant level, with Benjamini–Hochberg correction across all 10 pairwise comparisons.

### Software reproducibility and research use

The analysis used NeuroImgAgent 0.1.0, which requires Python 3.10 or later. External neuroimaging tools were integrated through tool manifests and configuration overlays. The prompts used for profiling, planning, tool routing, QC, failure diagnosis, reporting and consistency checking are provided in **Supplementary Note 1**. Large language models served as functional components of the evaluated system; during manuscript revision they were additionally used, under author supervision, for language editing and evidence organization. Model outputs did not replace any reported numerical analysis, and the authors verified all quantitative claims and citations in the manuscript.

NeuroImgAgent is intended for research use only and is not a medical device. QC verdicts, derived measures and generated reports require review by qualified researchers before publication or downstream interpretation.

## Data availability

Public datasets analysed in this study are available from OpenNeuro, CoRR, the Human Connectome Project, PRIME-DE and AFNI resources. Dataset identifiers, versions and access links are provided in **Supplementary Table 1**. The institutional UMR-5T human MRI dataset and awake-mouse MRI dataset are available on OpenNeuro under accession numbers ds008970 and ds009028, respectively, with version-specific DOIs 10.18112/openneuro.ds008970.v1.1.0 and 10.18112/openneuro.ds009028.v1.0.0. Source data underlying the figures are provided with this paper.

## Code availability

Development code is maintained at https://github.com/NeuroImgLab/NeuroImgAgent.

## Acknowledgements

We thank the investigators and data-sharing initiatives that made the neuroimaging datasets used in this study publicly available, and the developers and maintainers of the open-source software on which NeuroImgAgent builds. Data were provided in part by the Human Connectome Project, WU-Minn Consortium (Principal Investigators: David Van Essen and Kamil Ugurbil; 1U54MH091657) funded by the 16 NIH Institutes and Centers that support the NIH Blueprint for Neuroscience Research; and by the McDonnell Center for Systems Neuroscience at Washington University. We thank the investigators who contributed the University of Oxford macaque dataset to PRIME-DE. The acquisition and sharing of the Oxford data were supported by the Wellcome Trust, the Royal Society, the Medical Research Council UK and the Biotechnology and Biological Sciences Research Council UK. We thank Changjun Tie, Xiaoling Luo, Yingtong Wu and other staff members of the Paul C. Lauterbur Research Center for Biomedical Imaging (SIAT) as well as Lifei Huang of United Imaging Life Science Instrument, Wuhan, China, for their support and assistance.

## CRediT authorship contribution statement

**Jianyu Yuan**: Methodology, Software, Visualization, Validation, Investigation, Formal analysis, Writing-original draft, Writing-review & editing. **Hongtao Zhang**: Methodology, Software, Visualization, Investigation, Writing-original draft. **Liwen Wan:** Investigation, Resources, Writing-review & editing. **Yuanxi Shen**: Methodology, Software, Formal analysis, Investigation, Validation, Visualization. **Zhifei Zhu**: Methodology, Investigation, Software, Visualization. **Yexin Han**: Methodology, Software, Investigation. **Zeping Xie**: Visualization, Data curation, Writing-review & editing. **Pingfu Wang**: Investigation, Data curation, Writing-review & editing. **Qian Wan**: Methodology, Resources, Writing-review & editing. **Xiaojing Long**: Methodology, Resources, Writing-review & editing. **Chao Zou**: Methodology, Resources, Writing-review & editing. **Dong Liang**: Conceptualization, Funding acquisition, Investigation, Writing-review & editing. **Hairong Zheng**: Conceptualization, Funding acquisition, Investigation, Writing-review & editing. **Ye Li**: Conceptualization, Funding acquisition, Investigation, Writing-review & editing. **Yi He**: Conceptualization, Funding acquisition, Resources, Investigation, Visualization, Methodology, Software, Writing-review & editing, Supervision.

