## Supplementary Figures for "NeuroImgAgent: a multi-agent platform for cross-species fMRI analysis with evidence-gated quality control"

### Supplementary Information

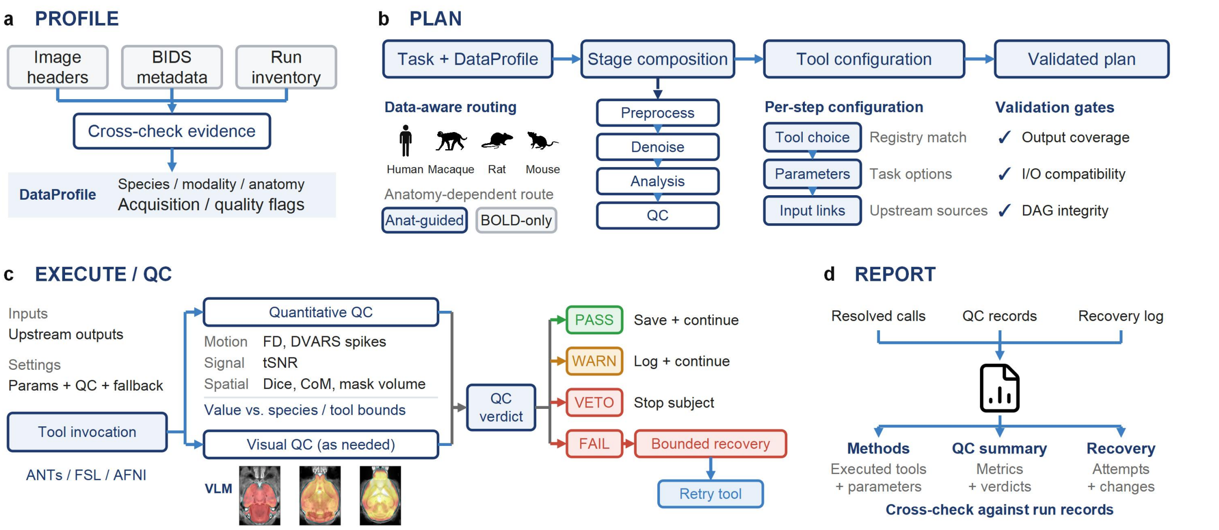


**Extended Data Fig. 1. Core module implementation details. a,** PROFILE combines image headers, Brain Imaging Data Structure (BIDS) metadata and run inventory to derive a DataProfile containing species, modality, anatomical-image availability, acquisition parameters and quality flags. **b**, PLAN uses the task and DataProfile to select species- and anatomy-appropriate routes, compose template-informed stages and configure tools, parameters and upstream inputs. Validation checks task-output coverage, input/output compatibility and directed acyclic graph integrity. Check marks denote validation gates rather than measured success rates. **c**, EXECUTE resolves inputs and settings before tool invocation. Quantitative QC assesses applicable motion, signal and spatial measures, including framewise displacement (FD), DVARS spikes, temporal signal-to-noise ratio (tSNR), Dice overlap, centre-of-mass (CoM) distance and mask volume. Vision–language model (VLM) inspection is used when required by the stage and policy. PASS and WARN permit continuation with records; VETO terminates subject processing; eligible FAIL outcomes enter bounded recovery and retry. Recovery depends on policy and does not guarantee success. The images illustrate existing mouse registration-QC evidence. **d**, REPORT combines resolved tool calls, QC records and recovery logs to generate Methods text, a QC summary and recovery history, with consistency checks against execution records. The figure depicts implementation mechanisms, not experimental validation of every route or automatic recovery in every case.

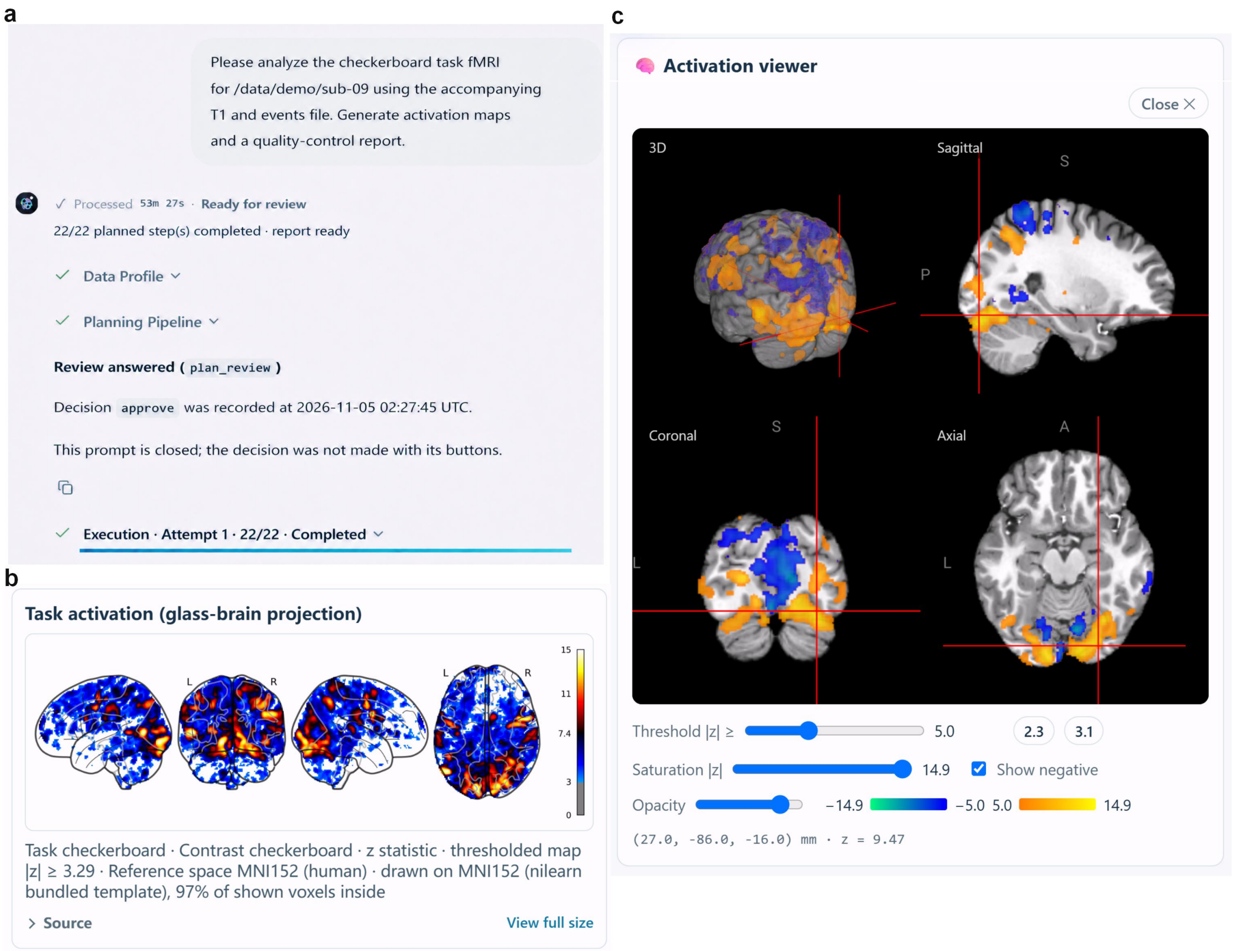


**Extended Data Fig. 2. Demonstration of task-based fMRI analysis using NeuroImgAgent. a,** The NeuroImgAgent user interface for submitting a task-based fMRI analysis request. **b,** Representative results displayed in the NIA web interface for a human subject performing a checkerboard task. Glass-brain projections of the thresholded z-statistic map in MNI152 space. **c,** The interactive activation viewer, with the statistical map overlaid on an anatomical image in three-dimensional, sagittal, coronal and axial views. Display controls allow adjustment of the threshold, colour saturation and opacity.

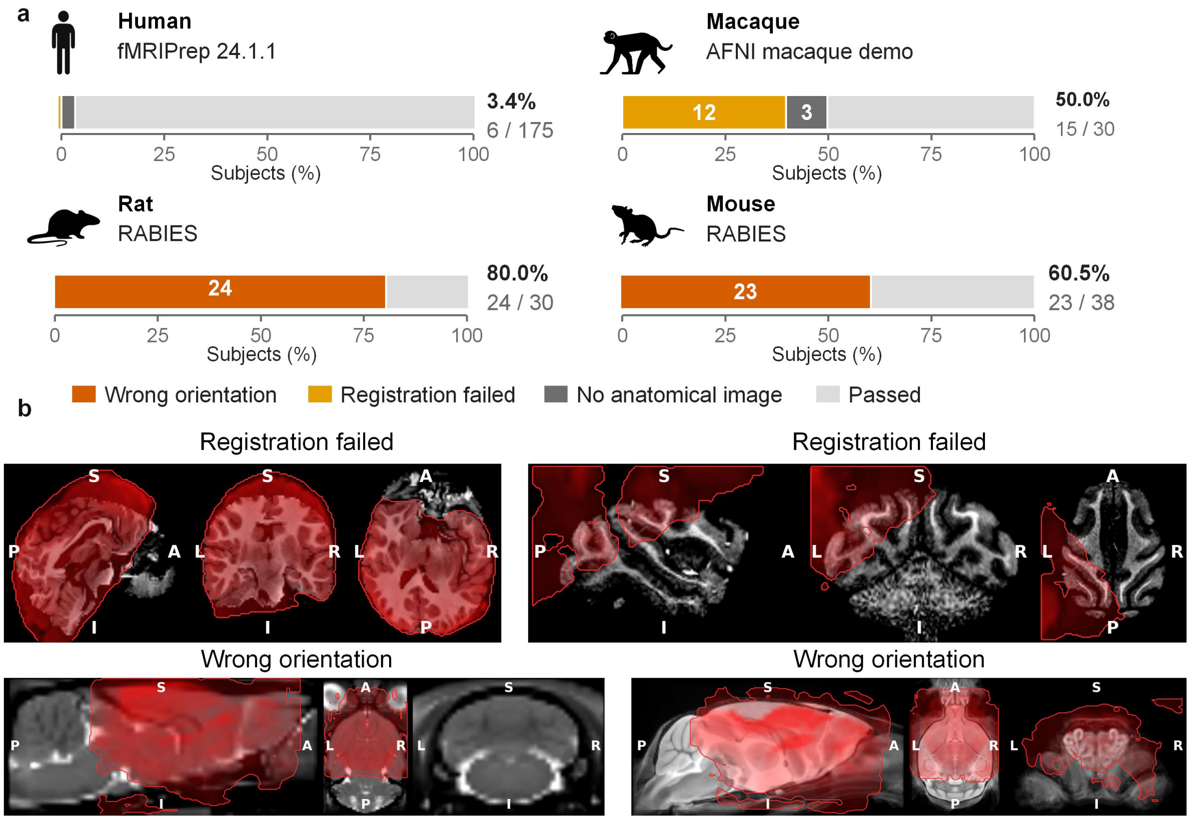


**Extended Data Fig. 3. Input incompatibilities and processing failures in baseline workflows without manual data preparation.** Each species' standard pipeline was run as distributed, without intervention: fMRIPrep 24.1.1 (human, n = 175), the AFNI macaque demo pipeline (macaque, n = 30) and RABIES 0.5.1 (rat, n = 30; mouse, n = 38). **a**, Outcome per subject, with failures of three types. No anatomical image: the pipeline cannot run. Registration failed: EPI/anatomical Dice below 0.6 (macaque), or 0.80, the lowest of 170 processed subjects (human). Wrong orientation: the native-space EPI differs from a manually re-oriented rerun by an axis flip or exchange; the 8 awake mice were judged from the raw image header, which labels coronal slices as axial. Left–right-only differences (6 rats) were not counted, as image content cannot confirm them. **b**, Representative examples of standard-pipeline failures per species. Human: the EPI is centred 12 mm posterior, leaving the frontal pole uncovered. Macaque: the EPI sits at the posterior–superior–left edge, severely misaligned. Rat and mouse: front–back reversal shifts the EPI anteriorly.

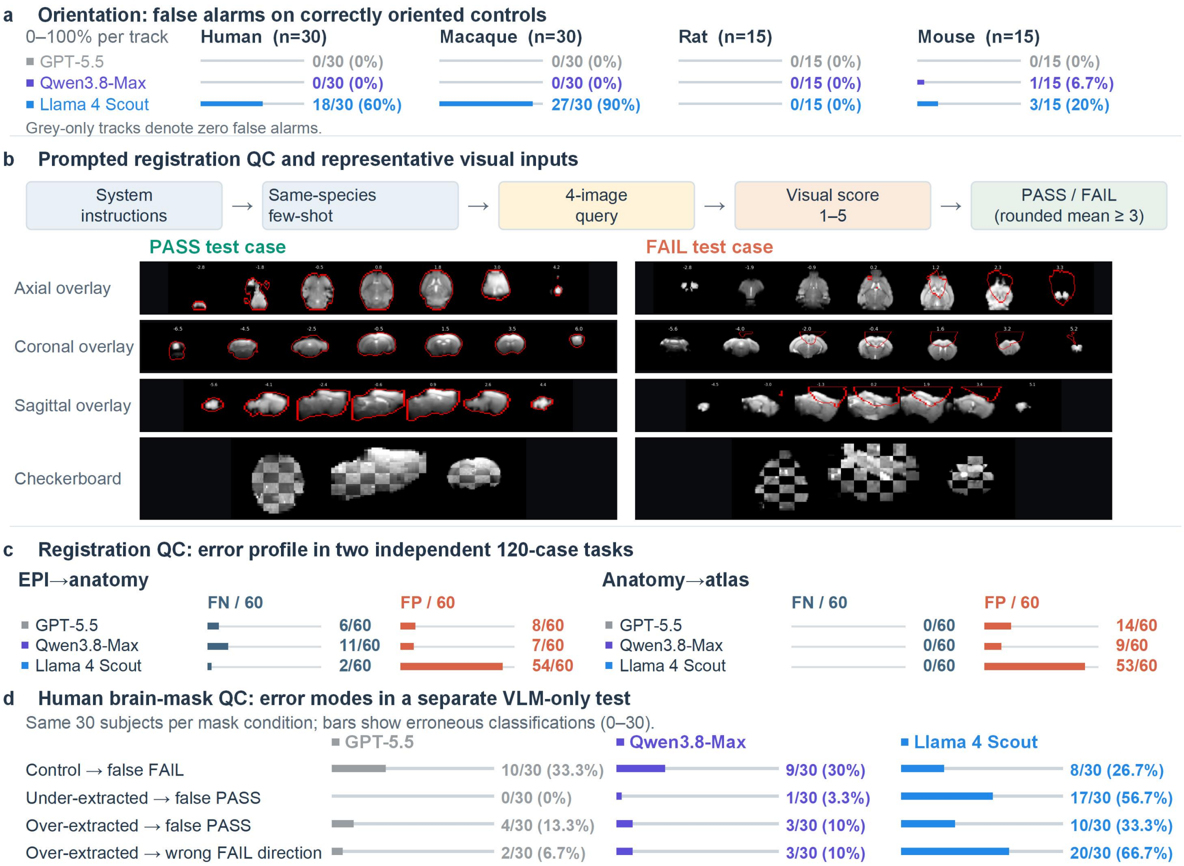


**Extended Data Fig. 4. Visual evidence and error patterns in VLM quality control. a**, False-alarm rates on correctly oriented controls (each track spans 0–100%); labels give count/evaluated controls (percentage) for all 12 model-by-species results. **b**, Prompt structure for registration QC and two representative test cases. Each query followed the system instructions and a same-species PASS/FAIL reference pair from held-out subjects (few-shot priors), and comprised four images of the test case: axial, coronal and sagittal overlays and a checkerboard image. The VLM rated five criteria on a 1–5 scale; the mean score was rounded half-up, and a value of at least 3 was classified as PASS. The two examples are scored mouse EPI-to-anatomy test cases. **c**, EPI-to-anatomy and anatomy-to-atlas each include 120 cases (30 per species; 15 true PASS and 15 true FAIL per species). FN counts true PASS cases called FAIL; FP counts true FAIL cases called PASS. Both bars and labels use a denominator of 60 per task and model. **d**, A separate VLM-only test on the same 30 human subjects per mask condition shows four error modes; labels give count/30 (percentage). A wrong FAIL direction means an over-extracted mask was called under-extracted.

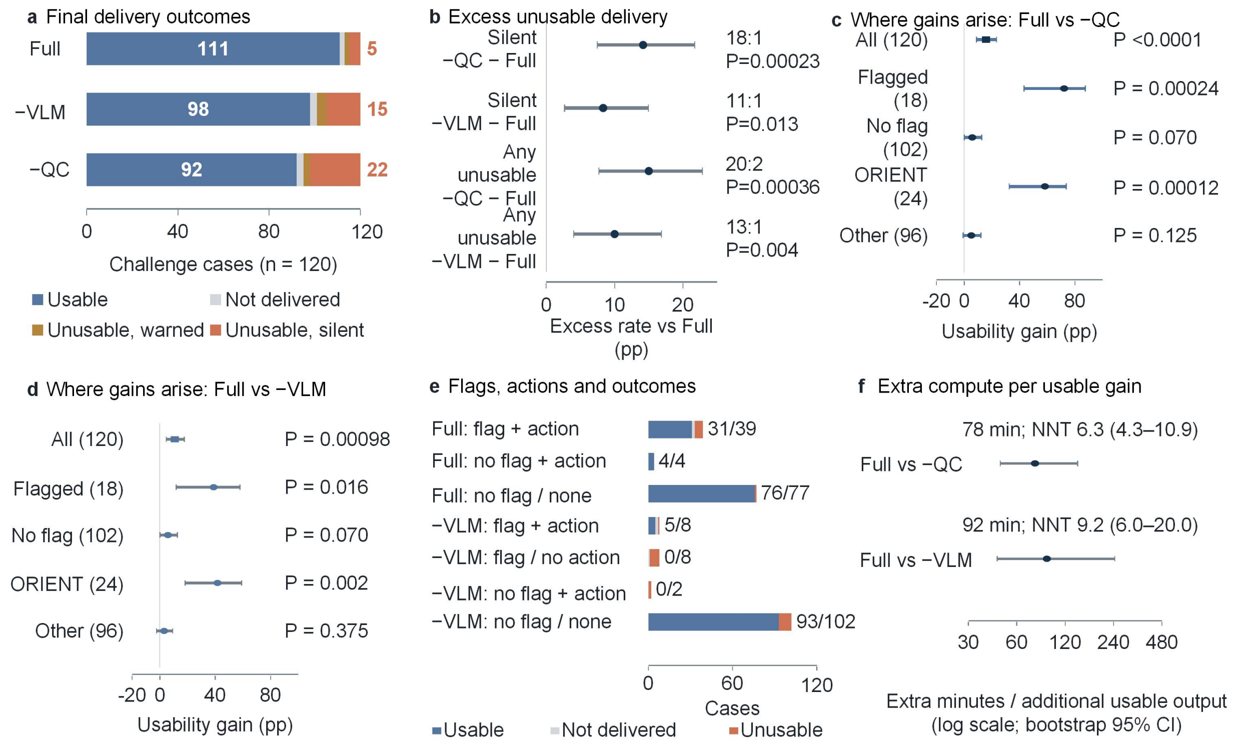


**Extended Data Fig. 5. Delivery, corrective actions and computational cost in the challenge cohort. a,** Final outcomes across 120 paired challenge cases. Not delivered includes deliberate stops and runs lacking outputs. **b,** Excess rates of silent or any unusable delivery in ablated arms relative to Full; 95% confidence intervals (CI), paired Newcombe method; P values, two-sided exact McNemar tests with Holm correction across three comparisons per endpoint. **c,d,** Full-minus-ablation success-rate differences by −QC metric flag or challenge type, with paired Newcombe 95% CI; displayed P values are unadjusted exact McNemar tests. The flagged stratum includes three no-output −QC runs. **e,** Recorded flag/action/outcome paths; taking action does not establish successful repair or temporal precedence of the flag. **f,** Extra compute per additional usable output (114 cases with delivery in all arms; species-stratified bootstrap 95% CI). NNT is the reciprocal of the net success-rate difference across all 120 cases. Cross-arm gains do not establish within-run causal repair or successful VLM reasoning; runtime estimates were not controlled for hardware or concurrency.

75 tasks across four capability levels and three models

✓ Pass  × Fail

**a L1 Intent recognition** — 28 tasks

| Scenario | | GPT-5.5 Human | GPT-5.5 Mouse | GPT-5.5 Macaque | GPT-5.5 Rat | Qwen3.8-Max Human | Qwen3.8-Max Mouse | Qwen3.8-Max Macaque | Qwen3.8-Max Rat | Llama-4-Scout Human | Llama-4-Scout Mouse | Llama-4-Scout Macaque | Llama-4-Scout Rat |
|---|---|---|---|---|---|---|---|---|---|---|---|---|---|
| S1 | Explicit resting-state intent | ✓ | ✓ | ✓ | ✓ | ✓ | ✓ | ✓ | ✓ | ✓ | ✓ | ✓ | ✓ |
| S2 | Explicit task-based intent | ✓ | ✓ | ✓ | ✓ | ✓ | ✓ | ✓ | ✓ | ✓ | ✓ | ✓ | ✓ |
| S3 | Implicit intent (language model) | ✓ | ✓ | ✓ | ✓ | ✓ | ✓ | ✓ | ✓ | ✓ | × | ✓ | ✓ |
| S4 | Product keyword | ✓ | ✓ | ✓ | ✓ | ✓ | ✓ | ✓ | ✓ | ✓ | ✓ | ✓ | ✓ |
| S5 | Species-label injection | ✓ | ✓ | ✓ | ✓ | ✓ | ✓ | ✓ | ✓ | ✓ | ✓ | ✓ | ✓ |
| S6 | Multi-signal inference | ✓ | ✓ | ✓ | ✓ | ✓ | ✓ | ✓ | ✓ | ✓ | ✓ | ✓ | ✓ |
| S7 | No anatomical image | ✓ | ✓ | ✓ | ✓ | ✓ | ✓ | ✓ | ✓ | ✓ | ✓ | ✓ | ✓ |
| Passed / total tasks | | 28/28 | | | | 28/28 | | | | 27/28 | | | |

**b L2 Planning** — 24 tasks

| Task | Task detail | GPT-5.5 | Qwen3.8-Max | Llama-4-Scout |
|---|---|---|---|---|
| L2-01 Human | Template: resting state | ✓ | ✓ | ✓ |
| L2-01 Mouse | Template: resting state | ✓ | ✓ | ✓ |
| L2-01 Macaque | Template: resting state | ✓ | ✓ | ✓ |
| L2-01 Rat | Template: resting state | ✓ | ✓ | ✓ |
| L2-02 Human | Template: task based | ✓ | ✓ | ✓ |
| L2-02 Mouse | Template: task based | ✓ | ✓ | ✓ |
| L2-02 Macaque | Template: task based | ✓ | ✓ | ✓ |
| L2-02 Rat | Template: task based | ✓ | ✓ | ✓ |
| L2-03 Mouse | Template: no anatomy | ✓ | ✓ | ✓ |
| L2-04 Human | Preprocessing only | ✓ | ✓ | ✓ |
| L2-04 Mouse | Preprocessing only | ✓ | ✓ | ✓ |
| L2-04 Macaque | Preprocessing only | ✓ | ✓ | ✓ |
| L2-04 Rat | Preprocessing only | ✓ | ✓ | ✓ |
| L2-05 Human | Add slice-timing correction | ✓ | ✓ | ✓ |
| L2-06 Human | Multi-echo combination guard | ✓ | ✓ | ✓ |
| L2-07 Human | Seed-based functional connectivity | ✓ | ✓ | × |
| L2-08 Human | Temporal filtering / ALFF | ✓ | ✓ | ✓ |
| L2-09 Human | Fieldmap trigger | ✓ | ✓ | ✓ |
| L2-10 Human | No fieldmap available | ✓ | ✓ | ✓ |
| L2-11 Human | Open-ended novel intent | ✓ | ✓ | ✓ |
| L2-12 Human | Open-ended tool request | ✓ | ✓ | ✓ |
| L2-13 Mouse | Prune slice-timing step | ✓ | ✓ | ✓ |
| L2-14 Human | Plan-contract validation | ✓ | ✓ | ✓ |
| L2-15 Human | Plan-hash consistency across 10 repeats | ✓ | ✓ | ✓ |
| Passed / total tasks | | 24/24 | 24/24 | 23/24 |

**c L3 Fault recovery** — 9 human tasks

| Task | Task detail | GPT-5.5 | Qwen3.8-Max | Llama-4-Scout |
|---|---|---|---|---|
| L3-01 | Rule-hit parameter tuning | ✓ | ✓ | ✓ |
| L3-02 | Rule miss: language-model root-cause analysis | ✓ | ✓ | ✓ |
| L3-06 | Artifact-contract violation | ✓ | ✓ | × |
| L3-07 | Cohort isolation | ✓ | × | ✓ |
| L3-08 | Checkpoint resume | ✓ | ✓ | ✓ |
| L3-09A | Circuit breaker: open | ✓ | ✓ | ✓ |
| L3-09B | Circuit breaker: unmet | ✓ | ✓ | ✓ |
| L3-10 | Fallback-chain success | × | × | × |
| L3-11 | Fallback-chain termination | ✓ | × | ✓ |
| Passed / total tasks | | 8/9 | 6/9 | 7/9 |

**d L4 Tool construction** — 14 tasks

| Task | Task detail | GPT-5.5 | Qwen3.8-Max | Llama-4-Scout |
|---|---|---|---|---|
| L4-01 | Route: build missing capability | ✓ | ✓ | ✓ |
| L4-02 | Route: reuse existing tool | ✓ | ✓ | ✓ |
| L4-03 | Route: clarify seed definition | ✓ | ✓ | × |
| L4-04 | Route: reject unsafe request | ✓ | ✓ | ✓ |
| L4-05 | Generate: temporal signal-to-noise ratio | ✓ | × | × |
| L4-06 | Generate: DVARS with robust outlier detection | ✓ | × | × |
| L4-07 | Generate: fALFF | ✓ | × | × |
| L4-08 | Generate: T2* mapping | ✓ | × | × |
| L4-09 | Generate: tissue-grouped carpet plot | ✓ | × | × |
| L4-10 | Generate: multi-run quality-control montage | ✓ | × | × |
| L4-11 | End-to-end: build fALFF | × | × | × |
| L4-12 | End-to-end: reuse fALFF | × | × | × |
| L4-13 | Transfer: temporal signal-to-noise ratio across species | ✓ | × | ✓ |
| L4-14 | Robustness: DVARS edge cases | × | × | ✓ |
| Passed / total tasks | | 11/14 | 4/14 | 5/14 |

**Extended Data Fig. 6. Task-level benchmark outcomes.** Task-level verdicts for three models across 75 tasks (225 task–model observations). **a**, Intent recognition (L1; 28 tasks): seven scenarios across four species, covering explicit resting-state and task-based intent, implicit intent, product keywords, species-label injection, multi-signal inference and absent anatomical images. **b**, Planning (L2; 24 tasks): template selection, preprocessing-only plans, conditional processing steps, multi-echo safeguards, seed-based connectivity, filtering/ALFF, fieldmap availability, open-ended requests, plan-contract validation and plan-hash consistency across ten repeats. **c**, Fault recovery (L3; nine human tasks): rule-hit parameter tuning, rule-miss root-cause analysis, artifact-contract violations, cohort isolation, checkpoint resumption, circuit-breaker conditions and fallback-chain success or termination. **d**, Tool construction (L4; 14 tasks): routing to build, reuse, clarify or reject; generation of imaging-analysis and QC tools; end-to-end fALFF workflows; cross-species transfer; and DVARS edge cases. Check marks/crosses indicate PASS/FAIL; summaries show passed/total tasks.

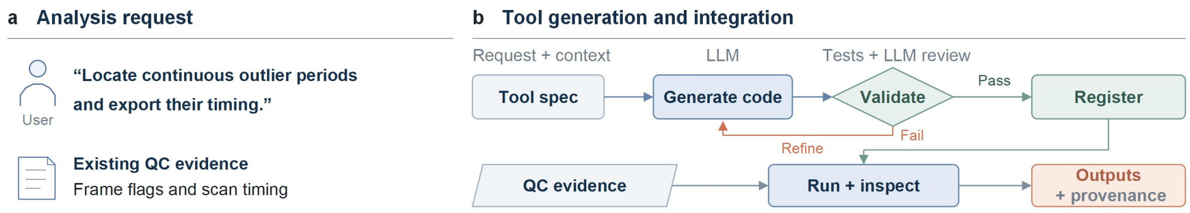


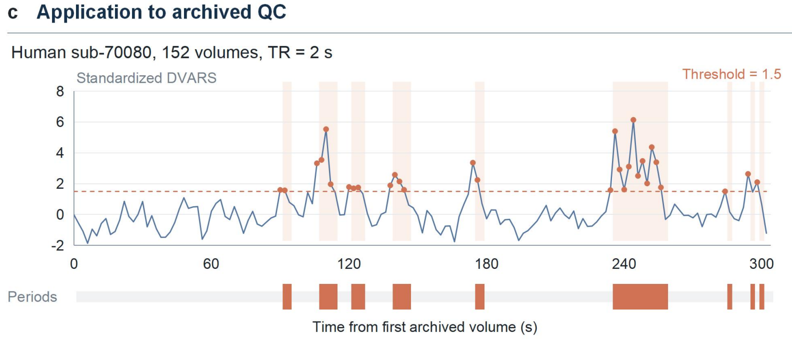


**Outlier periods**

| No. | Start (s) | End (s) | Δt (s) |
|---|---|---|---|
| 1 | 90 | 94 | 4 |
| 2 | 106 | 114 | 8 |
| 3 | 120 | 126 | 6 |
| 4 | 138 | 146 | 8 |
| 5 | 174 | 178 | 4 |
| 6 | 234 | 258 | 24 |
| 7 | 284 | 286 | 2 |
| 8 | 294 | 296 | 2 |
| 9 | 298 | 300 | 2 |

30 flagged frames, 9 periods, 60 s total

**Extended Data Fig. 7. Toolsmith workflow and an illustrative application to archived QC data.** **a,** A user requests continuous outlier periods using existing framewise QC evidence and scan timing. **b,** Schematic Toolsmith workflow: define the tool specification and input bindings, generate code, validate and review it, refine failed candidates, and register validated tools for execution and output inspection. Validation includes syntax, import and execution-contract checks and language-model code review. The diagram summarizes the implementation; it does not report a completed tool-generation experiment or human review. **c,** Offline interval extraction from an archived standardized DVARS trace for human sub-70080 (152 volumes; repetition time, 2 s). Frames with standardized DVARS >1.5 are flagged, and consecutive flagged frames are grouped into half-open intervals [start, end). Time zero is the first archived volume. Orange markers and shaded regions indicate flagged frames and derived intervals; the lower track and table report all nine intervals. Thirty frames are flagged, corresponding to 60 s in total. The complete trace is shown without smoothing or timepoint exclusion. The case illustrates a QC summary and does not estimate population performance, validate generated code or alter the original combined censoring policy.

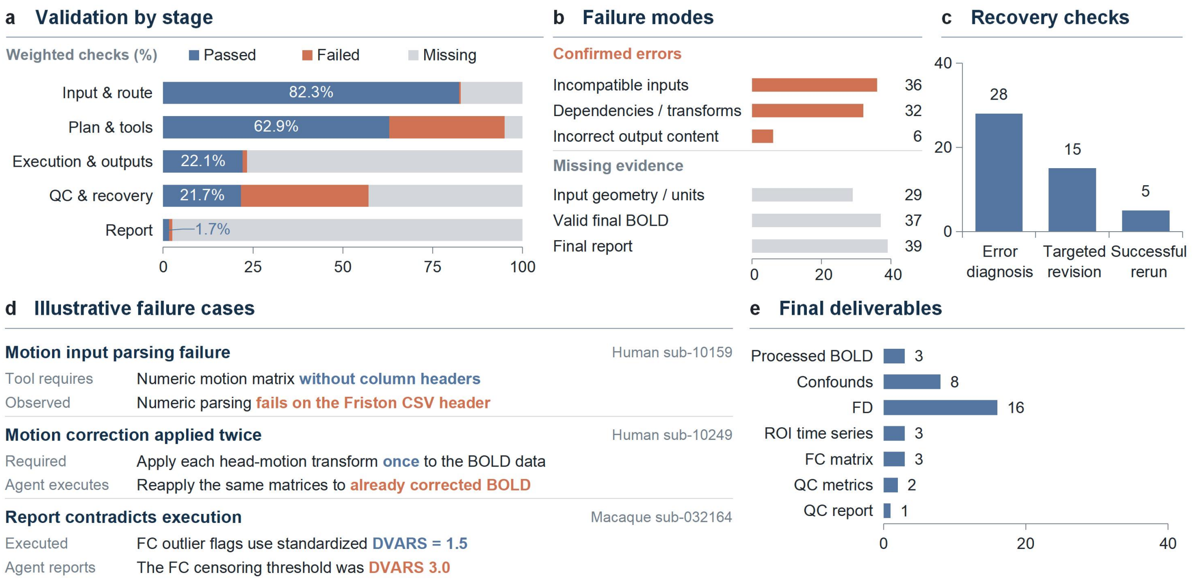


**Extended Data Fig. 8. Single-agent failure analysis.** Descriptive analysis of 40 single-agent development attempts across human, macaque, rat and mouse data (10 per species). **a**, Weighted validation outcomes by stage. Validation points within each stage, subjects and species receive equal weight. Passed, failed and missing-evidence fractions describe validation checks, not stage survival or end-to-end success rates. **b**, Numbers of attempts with non-exclusive confirmed-error or missing-evidence categories. An attempt may contribute to multiple categories. **c**, Recovery checks in the 38 attempts with recorded critical recoverable failures. Error diagnosis, targeted revision and successful rerun criteria were met in 28, 15 and five attempts, respectively. Two attempts without recorded critical failure chains are excluded. Counts describe attempt-level criteria across relevant failure chains, not individual errors or conditional transition probabilities. **d**, Illustrative recorded failures: parsing a headered motion-parameter CSV as a numeric matrix (human sub-10159); reapplying motion transforms to already corrected BOLD data (human sub-10249); and reporting a DVARS censoring threshold of 3.0 when functional connectivity used flags generated at standardized DVARS 1.5 (macaque sub-032164). **e**, Availability of final deliverables at termination, out of 40 attempts. A deliverable must have a current declared file binding, be readable and satisfy relevant axis-consistency checks. Counts do not establish scientific content correctness, complete QC or report factual accuracy; categories are non-exclusive. QC report denotes the HTML QC artifact. This development analysis does not establish a causal effect of agent architecture.

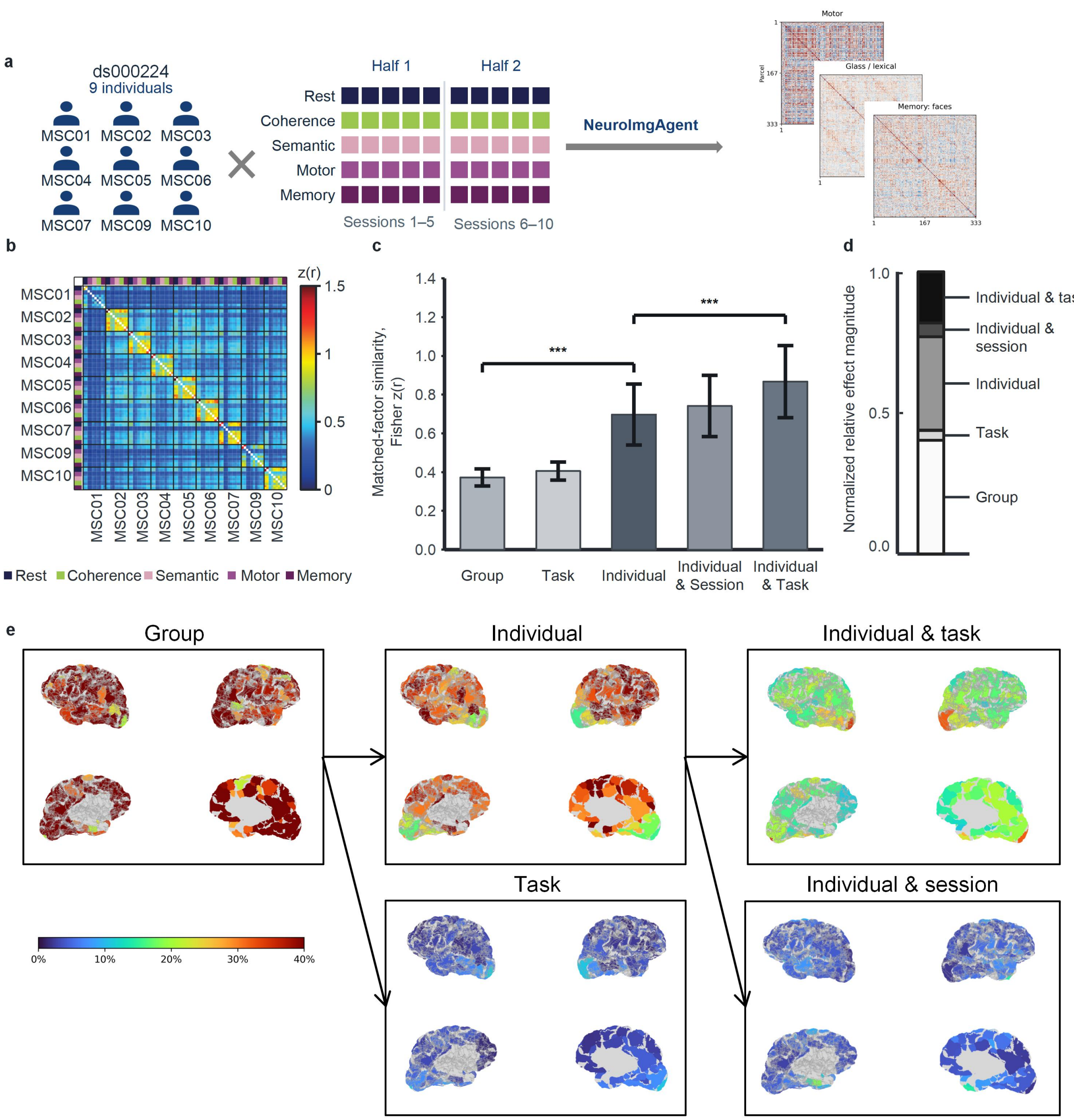


**Extended Data Fig. 9. Study design and decomposition of functional-network similarity. a**, Overview of the study design and analysis inclusion criteria. Scan data from nine participants (MSC01–MSC07, MSC09, and MSC10) in the public ds000224 dataset, acquired over 10 days and five states (rest and four tasks), were passed to NeuroImgAgent via natural language prompts for preprocessing and functional connectivity (FC) matrix generation. **b**, Pairwise functional-network similarity across 90 network instances from nine participants, comprising five states (rest, coherence, semantic, motor, and memory) and two independent data halves per state. Similarity is expressed as the Fisher-transformed correlation, z(r). White diagonal cells denote omitted self-comparisons, black boundaries separate participants, and the coloured strips indicate task states. **c**, Matched-factor similarity for the Group, Task, Individual, Individual × Session, and Individual × Task conditions. Bars represent the participant mean and error bars indicate ±1 SD. Brackets mark the contrasts from the original study that were replicated in the present dataset: Individual > Group and Individual × Task > Individual. Significance was assessed using two-sided paired t-tests with BH–FDR correction across 10 pairwise comparisons (***p < 0.001). **d**, Mean normalized relative contributions of the five components: Group, 40.2%; Task, 3.6%; Individual, 33.1%; Individual × Session, 4.9%; and Individual × Task, 18.1%. **e**, Cortical parcel maps of the normalized relative contributions of the Group, Individual, Individual × Task, Task, and Individual × Session components. Warmer colours indicate larger relative contributions; the colour scale ranges from 0% to 40%.
